# Hspa13 Regulates Endoplasmic Reticulum and Cytosolic Proteostasis Through Modulation of Protein Translocation

**DOI:** 10.1101/2022.06.27.497790

**Authors:** Mateo F. Espinoza, Khanh K. Nguyen, Melody M. Sycks, Ziqi Lyu, Maureen R. Montoya, Joseph C. Genereux

**Affiliations:** Graduate Program in Microbiology, University of California, Riverside, CA 92521; Department of Chemistry, University of California, Riverside, CA 92521

**Keywords:** Endoplasmic Reticulum, Import, Translocation, Proximity Labeling, Proteostasis, Molecular Chaperones, HSP70, Secretion

## Abstract

Most eukaryotic secretory proteins are co-translationally translocated through Sec61 into the endoplasmic reticulum (ER). Because these proteins have evolved to fold in the ER, their mistargeting is associated with toxicity. Genetic experiments have implicated the ER Hsp70 Hspa13/STCH as involved in processing of nascent secretory proteins. Herein, we evaluate the role of Hspa13 in protein import and the maintenance of cellular proteostasis. We find that Hspa13 interacts primarily with the Sec61 translocon and its associated factors. Hspa13 overexpression inhibits translocation of the secreted protein transthyretin (TTR), leading to accumulation and aggregation of immature TTR in the cytosol. ATPase inactive mutants of Hspa13 further inhibit translocation and maturation of secretory proteins. While Hspa13 overexpression inhibits cell growth and ER quality control, *HSPA13* knockout destabilizes proteostasis and increases sensitivity to ER disruption. Thus, we propose that Hspa13 regulates import through the translocon to maintain both ER and cytosolic protein homeostasis.

The raw mass spectrometry data associated with this manuscript has been deposited in the PRIDE archive and can be accessed at PXD033498.

## INTRODUCTION

Eukaryotic secretory proteins are primarily targeted to the endoplasmic reticulum (ER), where they fold, mature, and are trafficked to downstream environments (1). This targeting is achieved through N-terminal signal peptides or transmembrane domains (2). Proper targeting of secretory proteins to the ER promotes cellular fitness for two reasons. First, mistargeted protein deprives the organelle of its protein complement. Second, secretory proteins have evolved to mature and fold in the ER (3, 4). Signal peptidases, N-linked glycotransferases, disulfide isomerases, and other ER-localized proteostasis factors are absent from the cytosol. Without these factors, mistargeted secretory proteins cannot be properly processed and hence are prone to aggregation and toxicity. To ensure adequate quality control, multiple mechanisms and checkpoints promote targeting of signal peptide-containing proteins to the ER, primarily through engagement of the ribosome by the signal recognition particle (SRP) and subsequent co-translational translocation (5, 6). Some proteins, primarily those too short to engage SRP prior to synthesis, are delivered to the translocon through post-translational translocation (7, 8). Proteins persistently mistarget, for example due to weak signal sequences or cellular stress, are recognized by degradation pathways (9, 10).

The primary protein component of the eukaryotic translocon is Sec61, which is composed of three subunits (α1, β, γ) (11). A series of associated proteins further support protein translocation. For example, the primary ER Hsp70 BiP prevents protein misfolding during translocation, provides ratcheting to prevent backsliding, and is particularly important in assisting translocation for proteins harboring weak signal sequences (12, 13). The J-domain proteins Sec62 and Sec63 stimulate BiP ATPase function, and knockout studies suggest that they are particularly important for post-translational translocation (14–16). The OST complex promotes co- and post-translational transfer of high mannose glycans onto asparagine in N-linked glycosylation sequons (17, 18). Several literature reports over the past decade suggest that the poorly studied ER Hsp70 Hspa13/STCH might also be involved in protein import. First, Hspa13 localizes to the ER translocon Sec61, based on proximity labeling experiments (19). Hspa13 knockout sensitizes HCT116 human cells to the signal peptidase inhibitor cavinafungin, even more so than signal peptidase components, suggesting a role for Hspa13 in nascent protein maturation (20). Hspa13 expression is negatively associated with mouse lifespan following prion infection (21). Prion mislocalization to the cytosol, for example due to weak signal sequence, is associated with their pathogenicity (22–24). These findings were intriguing, given how little is known about Hspa13 function.

Hsp70 proteins are hub chaperones in the Hsp70 ATP-dependent chaperoning pathway and are present in most cellular compartments (25). As a chaperone, Hsp70 segregates unfolded proteins, preventing inappropriate interactions with other proteins while also promoting folding into the proper conformation (26– 28). Unlike other Hsp70s, Hspa13 lacks the substrate binding domain necessary for canonical Hsp70 functions (29, 30). (**Figure 1A**). The nucleotide binding domain has 39% identity with both BiP and the cytosolic Hsp70 HSPA1A, and Hspa13 is highly conserved across Eumetazoa (30, 31). Hspa13 is also a target of the Unfolded Protein Response (UPR), selectively induced by the XBP1s arm of the UPR (32, 33). It lacks a KDEL retention motif, leading to its stress-responsive secretion. Finally, reports have implicated Hspa13 in gastric cancer (34, 35), ER-associated degradation (ERAD) (36), plasma cell differentiation (37, 38), and inhibition of programmed cell death (39). However, none of these reports has identified a molecular role for Hspa13.

**Figure 1:**
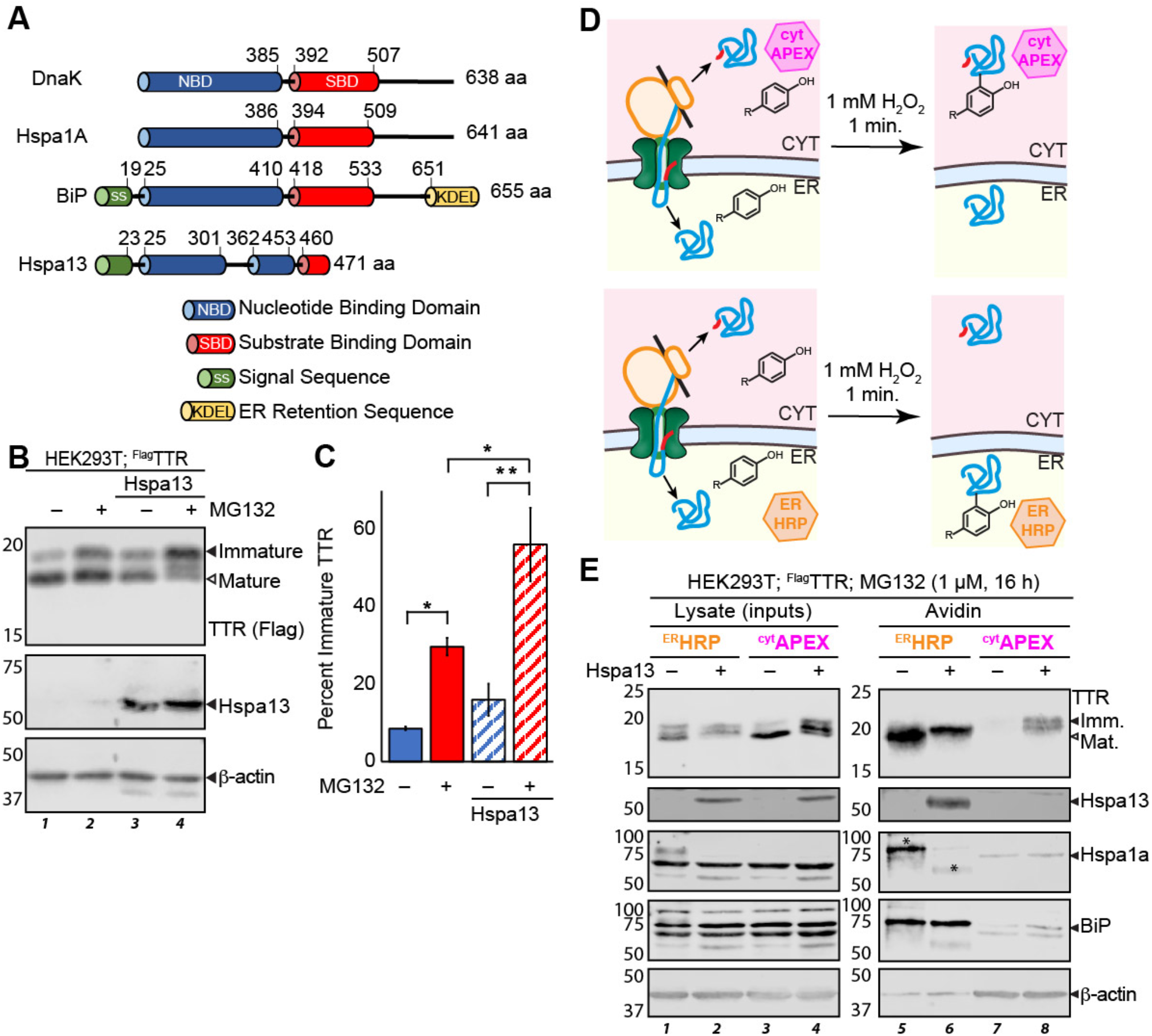
Hspa13 overexpression inhibits TTR import into the ER. **A**. Schematic comparing the domain structure of Hspa13 with those of the cytosolic human Hsp70 Hspa1a, the ER human Hsp70 BiP, and the *E. coli* Hsp70 DnaK. ATPase mutations are also highlighted. **B**. Representative immunoblot (n = 5) of SDS-PAGE separated lysates collected from HEK293T cells transiently co-overexpressing ^FLAG^TTR and Hspa13 as indicated. Cells were treated with either vehicle (0.1% DMSO) or 1 μM MG132 for 16 h prior to lysis. Molecular weight ladders are indicated on the left in kDa. **C**. Bar graph quantifying densitometry from **Fig 1B**. Percent immature TTR is determined as 100% x the ratio between the upper immature TTR band to the total monomeric TTR signal (n = 5; **, *p* < 0.01; *, *p* < 0.05). Signal ratios were analyzed using ANOVA (F = 15.44 > F_crit_ = 3.24) and then tested for significance with Tukey’s HSD test. Error bars represent standard error of the mean. **D**. Schematic illustrating the use of proximity labeling peroxidases to identify the ER and cytosolic proteomes. ^cyt^APEX is an engineered soybean ascorbate peroxidase, while HRP is an ER-directed horseradish peroxidase. **E**. Representative immunoblot of SDS-PAGE separated lysates from HEK293T cells overexpressing ^Flag^TTR and Hspa13 as indicated, and either ^ER^HRP or ^cyt^APEX. Cells were treated with biotin-phenol (BP; 30 min. pre-harvest) and H_2_O_2_ (1 min. pre-harvest), quenched, and the biotinylated protein purified from lysate using avidin beads. All cells were treated with 1 μM MG132 for 16 h prior to lysis. Hspa1a in lane 5 is obscured by interference from the TTR blotting step, marked with a *, as shown in **Figure S8**.

Herein, we evaluate whether Hspa13 affects secretory protein import into the ER in HEK239T cells. We find that Hspa13 overexpression inhibits translocation and that ATPase-inhibiting mutations in Hspa13 exacerbate this effect. Inhibited translocation decreases protein secretion, inhibits maturation for the population that still translocates, and leads to the accumulation of mistargeted and aggregated protein in the cytosol. Hence, we find that modulating Hspa13 abundance and activity leads to imbalances in cytosolic and secretory proteostasis.

## RESULTS

### Hspa13 overexpression inhibits TTR import into the ER

Transthyretin (TTR) is a homotetrameric secretory protein that is frequently used as a model substrate for ER protein homeostasis studies due to its extraordinarily well-characterized biophysical parameters, high secretory load, and disease relevance (40–42). TTR is particularly well-suited to ER import experiments, as a small population basally mistargets to the cytosol, this population increased with ER stress, and the immature and mature forms are readily separable by SDS-PAGE gel (43–46). To determine whether Hspa13 affects TTR localization, we co-overexpressed HEK293T cells with Hspa13 and Flag-tagged TTR (^Flag^TTR), with the Flag tag immediately C-terminal to the TTR signal peptide. Two TTR bands are apparent at 16 kDa and 18 kDa, consistent with the size of immature and mature ^Flag^TTR, respectively (**Figure 1B**). The upper band is smaller, implying that a majority of TTR present in the cell at steady-state has had the signal peptide cleaved. TTR contains a cryptic sequon that allows N-glycosylation of misfolded protein (47), however the bands fail to resolve upon treatment with the deglycosylase PNGase F (**Figure S1A**), indicating that the upper band represents immature ^Flag^TTR and not glycosylated protein. Hspa13 overexpression increases the abundance of immature ^Flag^TTR 1.9 ± 0.5 fold relative to the mature form (**Figure 1B,C**, lane 1 vs lane 3), consistent with ER import inhibition. Mistargeted secretory proteins are subject to degradation (9, 44). Cellular incubation with the proteasomal inhibitor MG132 for 16 hours increases total TTR both with and without Hspa13 overexpression (**Figure S1B**). In the presence of MG132, more immature TTR is observed, with Hspa13 overexpression still increasing immature ^Flag^TTR by 1.9 ± 0.2 fold. In addition, a small intermediate band appears between the mature and immature forms; we have not yet identified the nature of this species. In summary, if the immature band is indeed cytosolic, these results suggest that Hspa13 inhibits TTR import into the ER.

To better determine the localization of ^Flag^TTR proteoforms, we performed microsomal separation using digitonin, which selectively lyses the plasma membrane while leaving microsomes intact (**Figure S1C**). After microsomes are separated from the cytosol through precipitation, fractions are analyzed to determine protein distribution between two subcellular environments (48). We analyzed protein localization in HEK293T cells across a gradient of digitonin concentration. As expected, BiP is present in microsomal pellets (**Fig. S1D**) at all but the highest digitonin concentration. Overexpressed Hspa13 is also detectable in the pellet, confirming ER localization. However, the ^Flag^TTR monomers fractionate differently. While mature ^Flag^TTR co-fractionates with the ER proteins, immature ^Flag^TTR is preferentially found in the supernatant (**Fig. S1D**). Furthermore, while Hspa13 overexpression changes the relative amount of the two ^Flag^TTR populations (**Fig. S1D**), it does not affect their recovery with digitonin, indicating that Hspa13 impacts ^Flag^TTR localization in a manner dependent on signal peptide cleavage.

The digitonin assay can determine whether a homogenous population is a better fit between microsomal and cytosolic populations, but cannot as readily be used to differentiate whether a protein isoform is partitioned between two locations. Also, this assay copurifies cytosolic aggregates to the microsomal population (49). To better support cytosolic assignment for immature ^Flag^TTR, we applied our recently reported assay, which uses proximity labeling to measure ER protein mistargeting (46). Specifically, horseradish peroxidase with an N-terminal preimmunoglobulin signal sequence and C-terminal KDEL ER retention sequence (^ER^HRP) localizes to the ER (50), while an optimized ascorbate peroxidase (^cyt^APEX2) localizes to the cytosol (51). Cells are incubated with biotin-phenol, which distributes through the cell within 30 minutes (**Figure 1D**). Hydrogen peroxide is added, stimulating enzymatic production of biotin-phenoxy radicals that label endogenous proteins within a few nanometers of the peroxidases. Radical quenchers are added after one minute to minimize off-compartment labeling. Biotinylated proteins are enriched by avidin beads and quantified by Western blotting. The relative labeling of ER and secretory proteins by ^ER^HRP and ^cyt^APEX2 then indicates the extent of proper import as opposed to cytosolic mistargeting. As expected, ER resident proteins BiP and Hspa13 are preferentially labeled by ^ER^HRP, while cytosolic β-actin and Hspa1a are selectively labeled by ^cyt^APEX2 (**Figure 1E**). In the absence of Hspa13 overexpression, TTR is only labeled by ^ER^HRP and not by ^cyt^APEX2. Overexpression of Hspa13 increases the ^Flag^TTR that is labeled by ^cyt^APEX2. The migration of this cytosolic TTR is also consistent with that of the immature protein. Hence, Hspa13 inhibits ^Flag^TTR import and prevents its exposure to signal peptidase.

TTR is a known substrate of pre-emptive quality control, whereby activation of the UPR inhibits protein import into the cytosol (44). If Hspa13 overexpression activates the UPR, pre-emptive quality control could explain consequent ^Flag^TTR mistargeting. In contrast to Hyou1, whose overexpression robustly increases expression of the UPR target protein BiP, Hspa13 overexpression does not upregulate BiP, indicating that Hspa13 overexpression does not activate the UPR (**Figure S1E**). Another possibility is that Hspa13 inhibits signal peptide cleavage directly, with immature ^Flag^TTR then being rapidly retrotranslocated to the cytosol through ER-associated degradation (ERAD). However, we have previously demonstrated the inhibition of TTR signal peptide cleavage in the ER does not induce its cytosolic accumulation (46), but rather leads to ER accumulation of immature TTR instead.

### Hspa13 interacts with multiple components associated with the ER translocon

Hspa13 might inhibit ER TTR import through direct interactions with translocon components. To determine Hspa13 interactors, we employed affinity purification followed by mass spectrometry (AP-MS). We prepared Hspa13 with a C-terminal Flag (Hspa13^Flag^). We verified that Hspa13^Flag^ overexpression relocalizes ^Flag^TTR to the insoluble fraction similarly to untagged Hspa13 overexpression using ultracentrifugation (**Figure S2A**). Since regulatory interactions are often transient (52), we crosslinked cells with the cell-permeable crosslinker dithiobis(succinimidyl propionate) (DSP) prior to lysis and immunopurification. The crosslinking conditions were optimized to maximize recovery of the known Hspa13 and Sec61 interactor Bcap31 (37, 53) (**Figure S2B**). We found maximal Bcap31 recovery with 2 mM DSP treatment for 30 min., with only moderate background protein recovery from mock-transfected cells, as determined by silver stain (**Figure S2C**). Due to concerns over excessive protein loss, however, we limited DSP to 1 mM for the following experiment. Three plates of HEK293T cells were transfected with a mock plasmid (expressing eGFP) and three plates were transfected with Hspa13^Flag^. Two days post-transfection, cells were crosslinked, the crosslinking quenched, the cells lysed, the lysates normalized to total protein content, and those lysates immunoprecipitated over M2 anti-Flag agarose beads. The beads were stringently washed with RIPA buffer (50 mM Tris pH 7.4, 150 mM NaCl, 1% Triton X-100, 0.5% sodium deoxycholate, 0.1% SDS). Eluted proteins were processed for mass spectrometry, labeled with Tandem Mass Tags (TMT), and quantified by MuDPIT shotgun proteomics (**Figure 2A**). After filtering for common contaminants, 1298 proteins were identified. 1103 significant interactors were identified by using a previously reported method for correlating recovered protein levels to bait (Hspa13) levels (54), followed by significance testing with Storey’s modification of the Benjamini-Hochberg method to a 5% False Discovery Rate (55, 56) (**Figure 2B** and **Table S1**).

**Figure 2.**
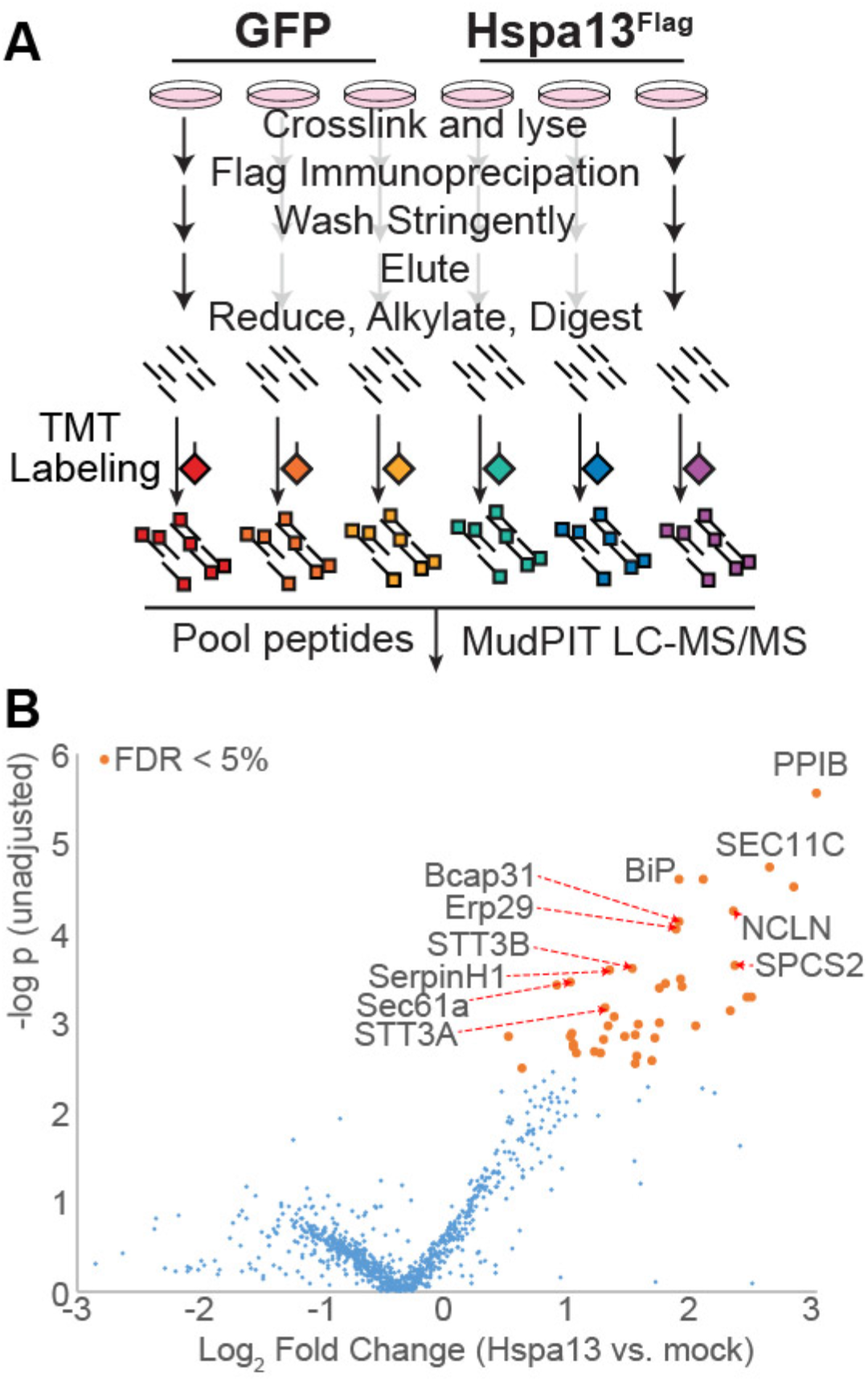
Hspa13 interacts with translocon-associated proteins. **A**. Schematic of immunoprecipitation and mass spectrometry to identify Hspa13 interactors. HEK293T cells overexpressing either mock (GFP) or Hspa13^Flag^ were crosslinked (1 mM DSP for 30 min), crosslinking quenched with 100 mM Tris pH, and the cells lysed. Lysates were anti-Flag immunoprecipitated and eluates processed for MS with tandem mass tag (TMT) labeling. **B**. Volcano plot for the relative co-IP recovery of proteins from HEK293T cells overexpressing Hspa13^Flag^ versus mock. Orange dots correspond to the set of proteins with q-values (see Methods for data analysis) below the False Discovery Rate threshold of 5%.

Hspa13 enrichment from Hspa13^Flag^ cells as compared to mock cells was 47.5 ± 4.1 fold, much lower than the theoretical ratio (∞) but consistent with the ratio compression often seen in TMT-MS2 experiments (57). These interactors include translocon components Sec61a1, Sec61b, NCLN, Sec62, and Sec63, as well as the signal particle recognition receptor component SRPRB and the translocon unclogger ZMPSTE24 (58, 59).

Four out of five components of the translocon-associated signal peptidase complex (SEC11C, SEC11A, SPCS2, and SPCS3) interact with Hspa13, as do seven of the twelve components of the oligosaccharyltransferase (OST) complex: RPN1, RPN2, DAD1, DDOST, MAGT1, STT3A, and STT3B (60). Although STT3B/MAGT1-containing OST complexes do not associate as strongly with the mammalian translocon, these complexes can still act co-translationally (17). We do not see association with members of the TRAP complex, which assists with translocation of transmembrane proteins and soluble proteins with a weak signal sequence (61–63). We do not see some reported cytosolic interactors, such as UBQLN2 (64, 65). It could be that our strong washing conditions prevent association with proteins that do not crosslink with Hspa13 in the ER. Another 11 interactors are major ER chaperones (i.e. BiP, ERdj3/DNAJB11, Grp94/HSPB90B1, HYOU1, Hsp47/Erp29, calnexin, PPIB, GANAB, PDIA3, PDIA4), many of which associate with nascent proteins to promote folding and prevent misfolding (66, 67). Several of these chaperones interact with BiP through direct interaction with the nucleotide binding domain (68, 69), which is similar to Hspa13. Hence, Hspa13’s interaction network is consistent with close association with the translocon and other components involved in maintaining proteostasis for nascent proteins.

### Mistargeted ^Flag^TTR is slowly degraded

Mistargeted secretory proteins can be degraded by the ubiquitin proteasome system (UPS) (4, 9, 10) or recruited into aggregates (70). We employed a cycloheximide (CHX) chase to characterize the degradation of ^Flag^TTR after Hspa13-induced mistargeting (**Figure 3A**). CHX arrests protein translation, so that existing populations of protein can be chased over time to determine their clearance kinetics. At the beginning of the chase, the relative proportion of immature protein is unaffected by the presence of MG132, suggesting that 2 h of proteasomal inhibition is not adequate to meaningfully affect the TTR population in the cell (**Figure 3A,B**). Hspa13 overexpression, on the other hand, nearly halves the relative mature fraction; MG132 treatment still has no further effect. Over the course of the chase, Hspa13 levels remain unchanged, suggesting a stable population. Mature ^Flag^TTR rapidly decreases in the presence or absence of Hspa13 (**Figure 3B** and **Figure S3**), consistent with its known secretion with a 3 h half-life (41, 71). The immature fraction, however, does not change over this timeframe, neither in the presence nor absence of MG132, indicating that its degradation is slow. After four hours the relative amount of mature and immature TTR plateaus to the same ratio under all conditions (**Figure 3C**). Hence, we can conclude that Hspa13-dependent mistargeted TTR accumulates and is persistent, with proteasomal degradation only becoming significant on the time-scale of days (**Figure 1C**). This relative resistance to proteasomal degradation could be due to the high thermodynamic and kinetic stability of TTR (40, 41), or to its propensity for aggregation if unable to fold (72).

**Figure 3.**
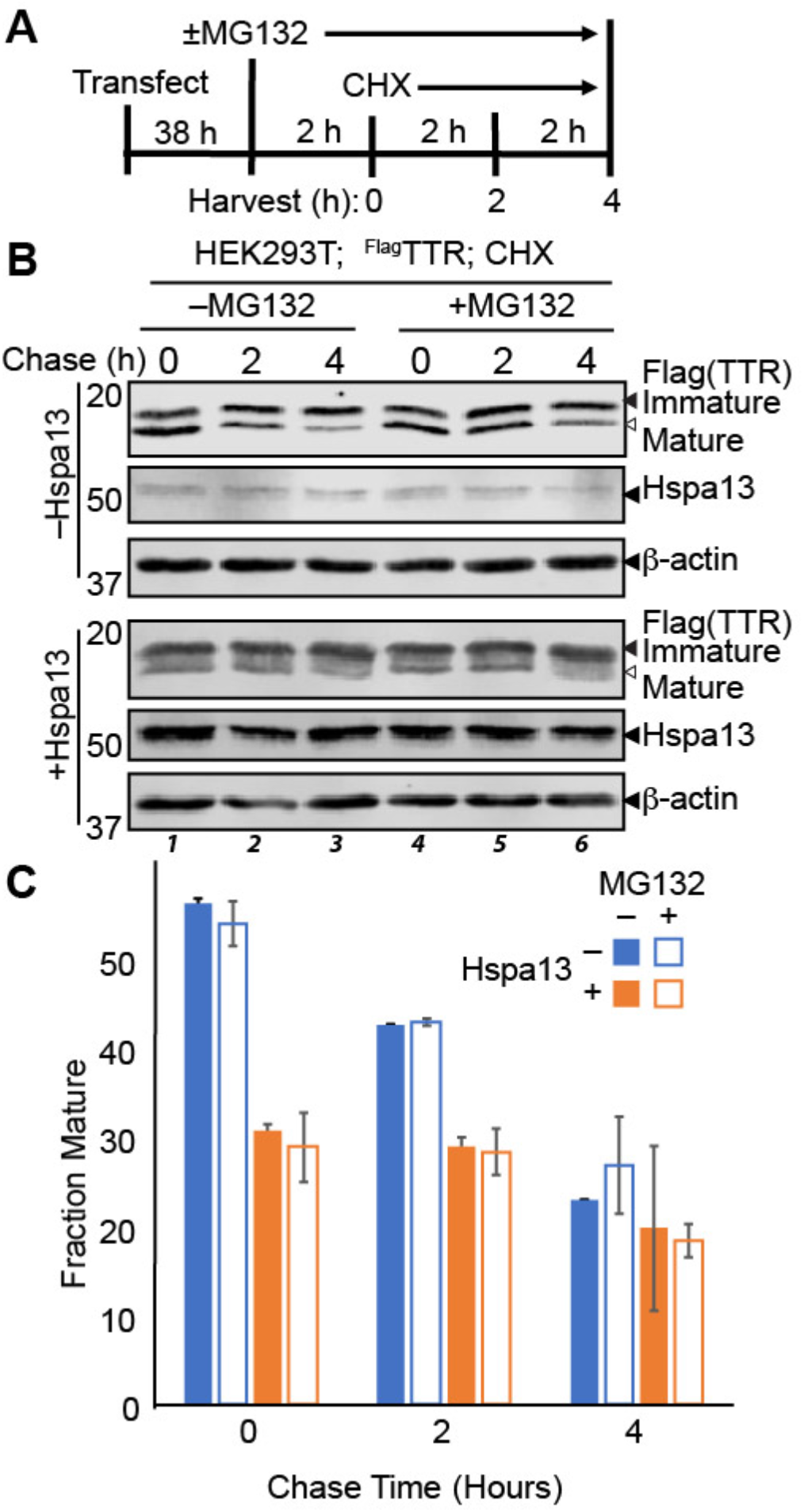
Immature TTR is slowly degraded. **A**. Schematic describing the timeline of the cycloheximide (CHX) chase experiment. Cells are treated with 10 μM MG132 or vehicle 2 h prior to the chase. The chase is initiated with 50 μg/mL CHX. Cells are harvested and lysed at indicated time-points. **B**. Representative immunoblot of SDS-PAGE separated lysates from HEK293T cells overexpressing ^Flag^TTR and Hspa13 treated as indicated. **C**. Densitometric quantifications of the fraction of mature TTR during the chase (n = 2). Error bars represent standard deviation.

### Mislocalized ^Flag^TTR forms insoluble aggregates

The accumulation of mislocalized protein presents a challenge for cytosolic protein homeostasis, as secretory proteins are prone to misfold and aggregate in the absence of proper maturation (3, 73). We hypothesized that accumulation of immature ^Flag^TTR in the cytosol following Hspa13 overexpression could lead to TTR aggregation. The thermodynamic stability of TTR amyloid aggregates makes them relatively resistant to denaturation, even when boiled in Laemmli buffer prior to SDS-PAGE gels (74, 75). Even native TTR tetramers tend to preserve some dimer structure under these conditions. To look for stable TTR aggregates, we co-overexpressed ^Flag^TTR and Hspa13 in HEK293T cells, limited boiling of lysates in reducing Laemmli buffer (including 2% SDS) to only 5 min., and separated lysates using gradient acrylamide SDS-PAGE gels that cannot effectively separate TTR isoforms but allow us to probe higher molecular weight ranges (**Figure 4A, B**). Expression of ^Flag^TTR alone leads to a prominent dimer band, but about half of the TTR is present in the monomer range. By contrast, Hspa13 overexpression leads to over 80% of TTR presenting as high molecular weight. Similar results were also obtained in SH-SY5Y cells (**Figures S4A**). Consistent with aggregates, long boiling times in Laemmli buffer reduce the high molecular weight fraction, with little aggregate remaining after 30 min. (**Figure S4B**). An alternative explanation for high molecule weight TTR immunoreactivity could be TTR ubiquitination. To test this hypothesis, we overexpressed C-terminally HA-tagged ubiquitin (Ub^HA^) (76) alongside TTR^Flag^ and Hspa13, and lysed in the presence of 12.5 mM N-ethylmaleimide to inactivate cellular deubiquitinases. The highly destabilized TTR^D18G^, a canonical ERAD substrate (77), was transfected as a positive control for a known ubiquitinated protein (78). As expected, anti-Flag immunoprecipitation of ^Flag^TTR^D18G^ provides a robust ubiquitination signature that is entirely dependent on proteasomal inhibition with MG132 (**Figure S4C**). The wild-type ^Flag^TTR, unexpectedly, shows noticeable ubiquitination even in the absence of MG132. This could be related to the slow clearance of mislocalized TTR (**Figure 3**). Nevertheless, Hspa13 does not affect the ubiquitination of TTR, demonstrating that high molecule weight TTR under conditions of Hspa13 co-overexpression is not due to increased ubiquitination.

**Figure 4.**
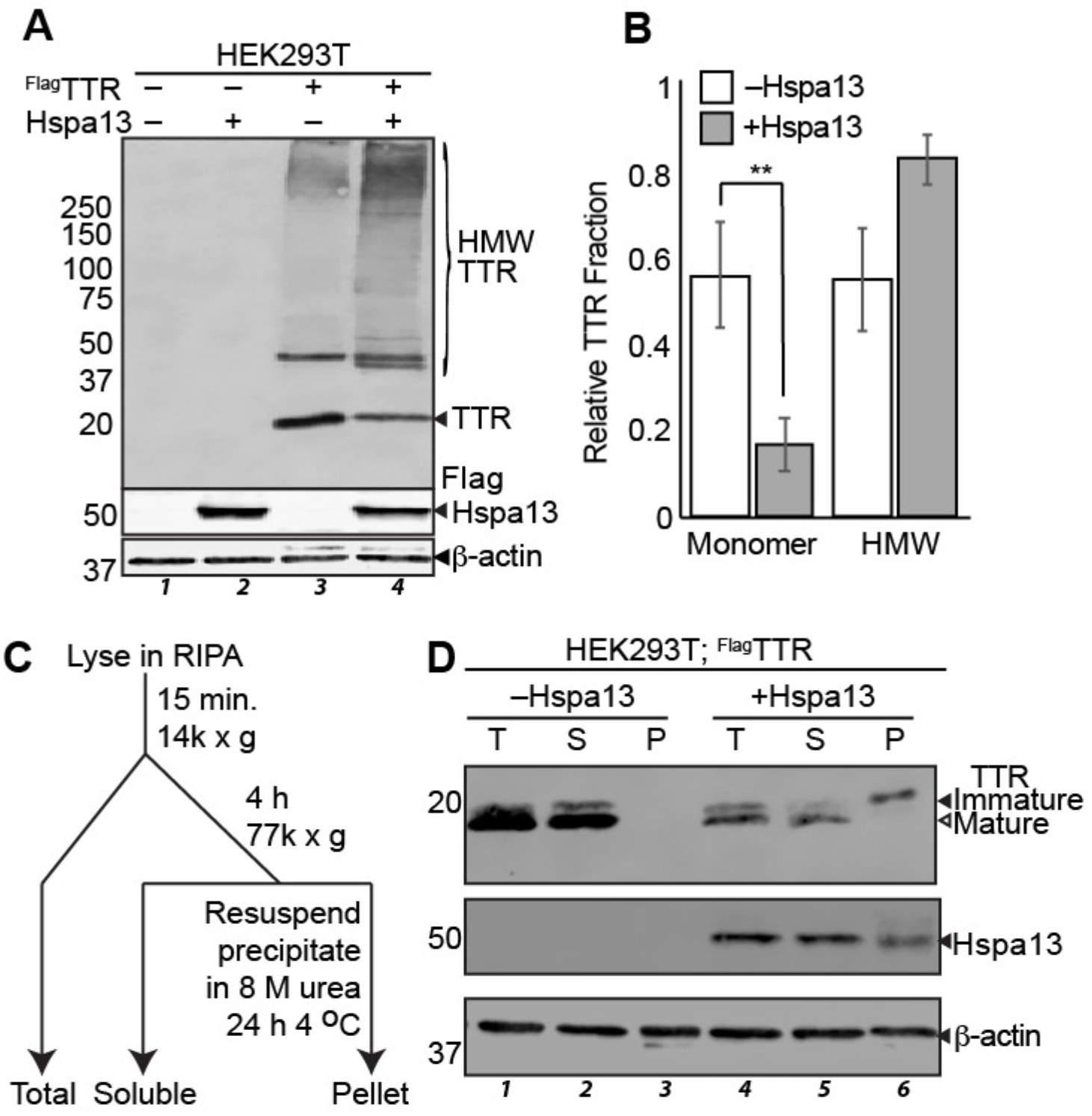
Mislocalized ^Flag^TTR forms insoluble aggregates. **A**. Representative immunoblot of lysates from HEK293T cells co-overexpressing ^Flag^TTR and Hspa13 as indicated. Lysates were boiled for 5 min. in Laemmli prior to separation by 4-20% gradient reducing SDS-PAGE. HMW indicated high molecular weight, as opposed to monomeric, TTR species. **B**. Densitometric quantification of monomeric and high molecular weight TTR fractions (n = 8). Error represents standard error of the mean. ** p < 0.01 by paired two-tailed Student’s t-test. **C**. Schematic describing ultracentrifugation to separate protein aggregates. **D**. Representative immunoblot of SDS-PAGE separated lysates from HEK293T cells overexpressing ^Flag^TTR and Hspa13 as indicated. Lysates were fractionated as described in panel **4C**. “T” indicates Total lysate before centrifugation; “S” indicates Soluble fraction; “P” indicates the Pellet. Equal percentages of pre-spin lysate were added to enable mass balance.

To further probe changes in TTR solubility with Hspa13 overexpression, we used ultracentrifugation to separate insoluble aggregates for cellular lysate (**Figure 4C**). After clearing chromosomal fractions and cellular debris from protein lysates with a 14,000 *x g* spin for 15 min., lysates were quantified, normalized for protein content, and then subjected to ultracentrifugation at 77,000 *x g* for 4 h at 4 °C. The resulting pellets were washed with RIPA buffer, and then solublized by incubating in 8 M urea for 24 h at 4 °C. In the absence of Hspa13 overexpression, ^Flag^TTR is completely soluble under these conditions (**Figure 4D**). Hspa13 overexpression induces a population of insoluble ^Flag^TTR, and the migration of this population is consistent with it being immature. These results support a model wherein Hspa13 overexpression inhibits ^Flag^TTR import, leading to the accumulation of immature, aggregation-prone TTR in the cytosol. We also compared the effect of Hspa13 overexpression to UPR induction by thapsigargin (Tg), which promotes secretion of aggregation-prone conformations of TTR (79). While Tg upregulates the UPR target BiP, and modestly inhibits TTR import efficiency (44, 46), it does not increase insoluble TTR to a similar extent as Hspa13 overexpression does (**Figure S4D**).

We considered that Hspa13 might have a similar effect on endogenous HEK293T proteins. To discover such proteins, we transfected HEK293T cells with either Hspa13 or mock (eGFP in the same plasmid vector). Total and insoluble lysate were prepared according to the same scheme of **Figure 4C**, and triplicates prepared for multiplexed quantitative proteomics with a single run for the six total lysate samples, and a single run for the six pelleted samples (**Figure S4E**). After filtering likely contaminants, we quantified 3372 proteins in the lysates and 1471 proteins in the pellets. We see no meaningful and significant difference in the cellular proteomes for cells overexpressing Hspa13 as compared to those expressing eGFP (**Figure S4F** and **Table S2**), with the exception of Hspa13 and GFP themselves. Crnkl1 (Fold change = 0.62 ± 0.21, p (moderated) = 8.7 × 10^−5^, qBH = 0.098) and Dimt1 (Fold change = 0.23 ± 0.08, p (moderated) = 7.6 × 10^−4^, qBH = 0.43) are possibly differential, but do not pass our threshold of qBH < 0.05. Similarly, in the pelleted insoluble fraction (**Figure S4G** and **Table S3**) only Hspa13 and GFP pass our significance threshold, though Fasn (Fold change = 1.17 ± 0.04, p (moderated) = 5.2 × 10^−4^, qBH = 0.26) and Myh10 (Fold change = 1.36 ± 0.16, p (moderated) = 3.7 × 10^−3^, qBH = 0.44) are potentially differential. Hence, under basal conditions, Hspa13 overexpression remodels neither the proteome nor its solubility.

### Hspa13 ATPase mutants exacerbate TTR import deficits and aggregation

Similar to other Hsp70 proteins, Hspa13 contains a nucleotide binding domain and is an active ATPase (29). In most Hsp70s, ATP hydrolysis is stimulated by client and Hsp40 binding, leading to a conformational change in the substrate-binding domain that increases affinity for and unfolding of the client protein (27). ATPase activity can also be increased with peptide sequences that mimic full-length clients (80). Unlike other Hsp70’s, Hspa13 ATPase activity is not stimulated by peptide binding, consistent with its lack of a substrate binding domain (29). In the absence of a clear function for Hspa13 ATPase activity, we considered that this activity might influence its effect on ER import. We prepared two Hspa13 constructs that we expected to be ATPase deficient, based on the high conservation of this domain between Hspa13 and other Hsp70s, and the well-characterized nature of these mutations across multiple Hsp70 proteins (**Figure 5A**). Briefly, BiP T229G and DnaK T199A do not affect ATP binding but decrease ATP hydrolysis by ≥ 95% (81, 82). Similarly, DnaK K70A and Hsc70 K71A bind ATP but cannot hydrolyze it (83, 84). We assessed Hspa13^K100A^ and Hspa13^T230A^ for their induction of immature and aggregated TTR using the ultracentrifugation assay (**Figure 5B**). As with the wild-type Hspa13, overexpression of either ATPase inactive Hspa13^K100A^ or Hspa13^T230A^ increases the amount of immature and insoluble TTR. This effect is greater for Hspa13K^100A^ than for the wild-type, and Hspa13^T230A^ expression partitions almost all TTR to the immature and insoluble population. ^Flag^TTR contains that Flag tag adjacent to the native TTR signal peptide cleavage site, which could affect behavior of the signal peptide. We hence also evaluated the effect of Hspa13 mutant co-expression with untagged TTR. Although Hspa13^K100A^ and Hspa13^T230A^ still inhibit TTR maturation and promote its aggregation, the effect of Hspa13^WT^ is muted (**Figure S5**), implying that the effect of Hspa13 on import is sensitive to signal peptide context. Although we cannot yet speculate as to the role of ATPase function regarding protein import into the ER, it is clear that the effect is exacerbated by loss of ATPase function. Impaired translocon function could be deleterious to cellular fitness. To test this hypothesis, we overexpressed Hspa13^WT^, Hspa13^T230A^, or eGFP (mock) in the same plasmid vector in HEK293T cells. Cellular proliferation was determined by the resazurin metabolic assay. Indeed, Hspa13^WT^ and Hspa13^T230A^ overexpression decrease cellular viability by 18% ± 1% and 36% ± 2% respectively (**Figure 5C**).

**Figure 5.**
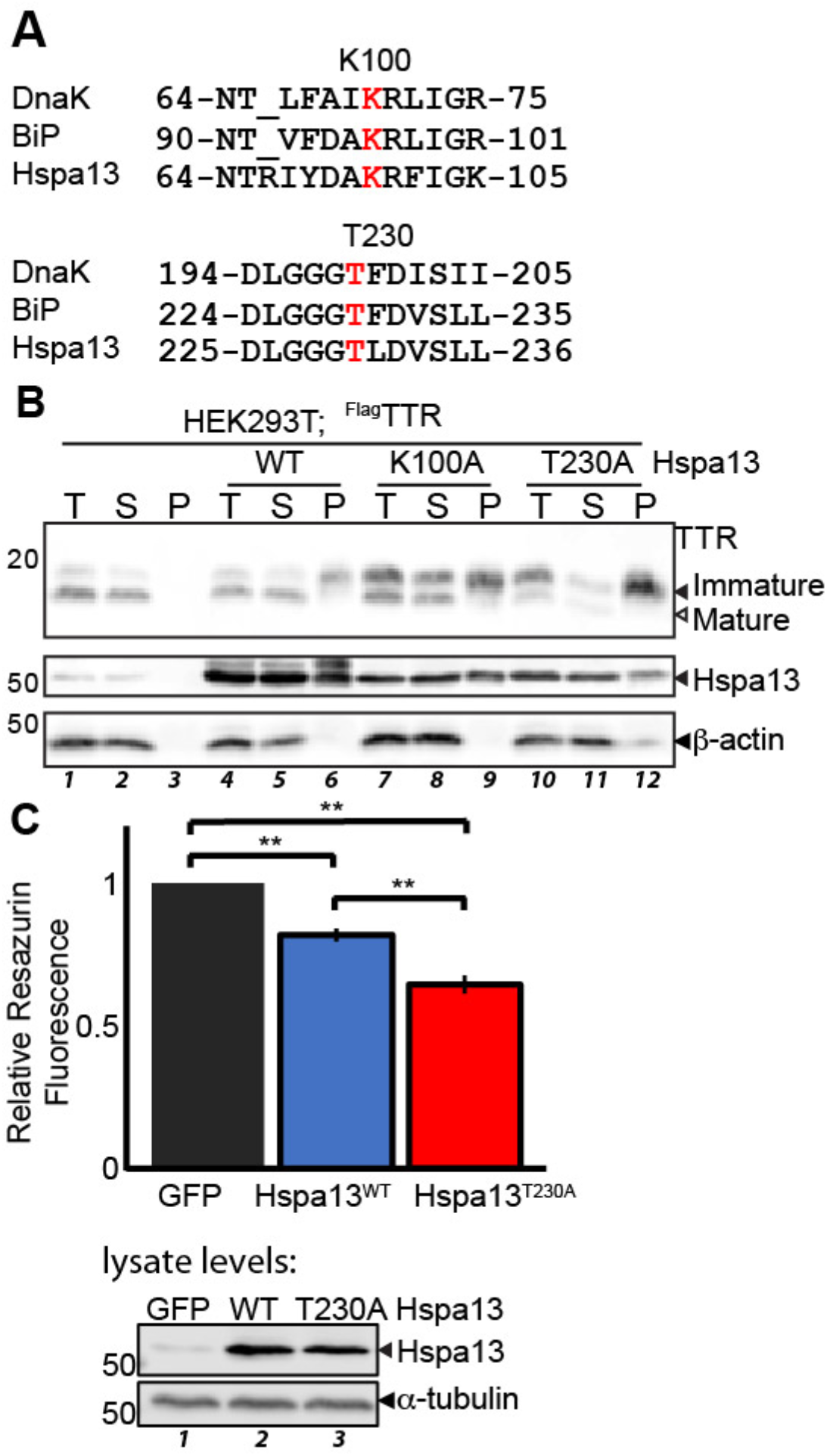
Hspa13 ATPase mutants exacerbate TTR import deficits and aggregation. **A**. Schematic depicting conservation between DnaK, BiP, and Hspa13 in the vicinity of known DnaK/BiP ATPase mutants. The sites of mutation are indicated in red. **B**. Representative immunoblot of SDS-PAGE separated lysates from HEK293T cells overexpressing ^Flag^TTR and Hspa13 as indicated. Lysates were fractionated as described in **Figure 4C**. “T” indicates Total; “S” indicates Soluble fraction; “P” indicates the Pellet. **C**. Normalized (to mock GFP overexpression) resazurin fluorescence (ex: 560 nm; em: 570 nm) from HEK293T cells overexpressing Hspa13 variants as indicated. Cells were seeded at 7,500 cells/well and wells individual transfected. The next day, and transfected with 21 uL of DNA/Ca_3_(PO_4_)_2_ for 16 h. Cells recovered with fresh media, then were incubated with media containing resazurin (25 μg/mL) for 2 h prior to measurement. Significance was assessed by ANOVA (n = 5; F = 182 > F_crit_ = 3.9) followed by Tukey’s post-hoc HSD. ** p < 1 × 10^−5^. An immunoblot from cells transfected in parallel is shown below.

### Hspa13 overexpression and mutation impair secretory proteostasis

Translocation kinetics are closely tied to processing and maturation in the ER. Signal peptide cleavage frees the C-terminus of lumenal proteins for co-translational processing and folding (85). Poorly optimized signal sequences demonstrate decreased glycan occupancy (86, 87). If Hspa13 modulation disrupts translocon function, that could also affect proteostasis of secretory proteins that still successfully enter the ER. Consistent with this hypothesis, Hspa13 co-overexpression inhibits NKCC2 glycan maturation in HEK293 cells (36). To evaluate proteostasis of mature TTR, we exploited the well-characterized affinity of ER chaperones for destabilized TTR (78, 88, 89). ^Flag^TTR was co-overexpressed with either Hspa13 or the other ER Hsp70, BiP. A C-terminal myc tag on BiP allows us to differentiate between the transfected and endogenous protein. ^Flag^TTR^D18G^, a highly destabilized variant of TTR, was also transfected as a positive control, and each sample immunoprecipitated to see TTR-associated chaperones. As previously reported, TTR^D18G^ robustly immunopurifies with both BiP and the ER Hsp40 ERdj3 (**Figure 6A**). ^Flag^TTR^WT^, which is both kinetically and thermodynamically stable, does not associate with either chaperone. BiP overexpression is sufficient to induce some BiP co-purification with TTR^WT^, but does not induce ERdj3 association. This suggests that while BiP overexpression can drive BiP association to TTR^WT^, it does not induce destablization of the protein. Hspa13 overexpression, however, induces binding of both BiP and ERdj3 to TTR^WT^, indicating a destabilized population of TTR in the ER. This suggests that ER proteostasis is impaired by Hspa13 overexpression. It is worth noting that BiP overexpression also leads to an increase in immature TTR (**Figure 6A**, lanes 3 and 7), and this immature population is localized to the cytosol (**Figure S6A**). However, this effect is not simply due to overexpression of abundant ER chaperones, as overexpression of ERjd3 does not affect TTR maturation and localization (**Figure S6A**). Similarly, HYOU1 overexpression only slightly affects TTR localization (**Figure S6A**), consistent with UPR activation by HYOU1 and consequent BiP overexpression (**Figure S1E**). A more detailed investigation of BiP overexpression and TTR translocation will be addressed in a separate manuscript.

**Figure 6.**
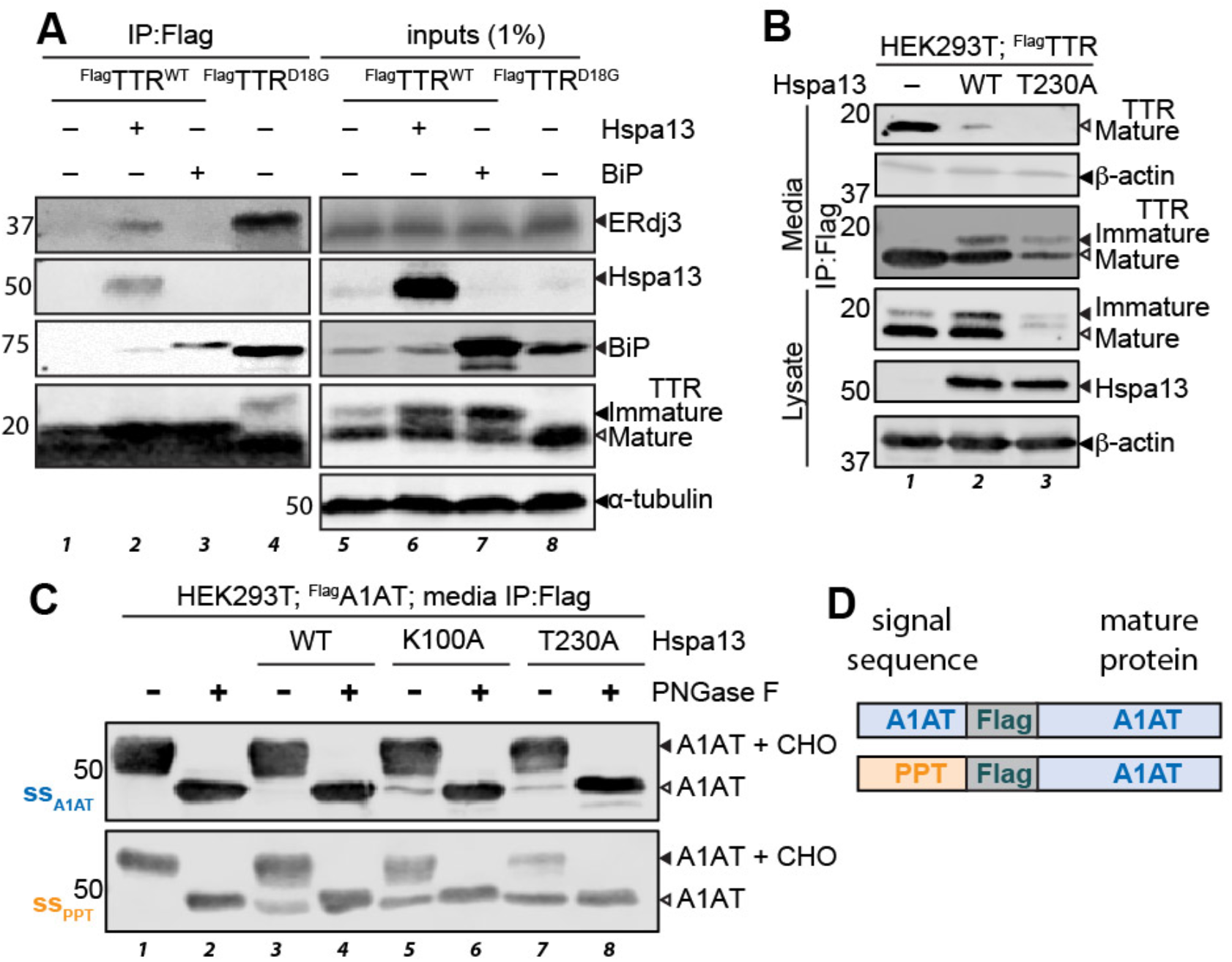
Hspa13 overexpression and mutation impair secretory proteostasis. **A**. Representative immunoblots of SDS-PAGE separated lysates and M2 anti-Flag immunoisolates from HEK293T cells overexpressing ^Flag^TTR variants, Hspa13, and hamster BiP^myc^ as indicated. **B**. Representative immunoblots of SDS-PAGE separated lysates, conditioned media, and conditioned media M2 anti-Flag immunoisolates from HEK293T cells overexpressing ^Flag^TTR and Hspa13 variants as indicated. Media were conditioned for 29 h. Quantification in **Figure S6B C**. Representative immunoblots of SDS-PAGE separated M2 anti-Flag immunoisolates from HEK293T cells overexpressing ss_A1AT_^Flag^A1AT or ss_PPT_^Flag^A1AT and Hspa13 variants as indicated. Media were conditioned for 24 hours prior to IP. **D**. Schematic illustrating the two A1AT constructs employed.

Since TTR ER proteostasis is inhibited by Hspa13 overexpression, we considered the effect of Hspa13 overexpression on secretion. HEK293T cells were transfected with ^Flag^TTR and either Hspa13^WT^ or Hspa13^T230A^, the media changed after 16 h, and then media conditioned for another 24 h. Overexpression of Hspa13^WT^ nearly eliminates ^Flag^TTR secretion, despite a substantial mature population persisting in the cells (**Figure 6B** and **Figure S6B**). Following Hspa13^T230A^ co-overexpression, almost no ^Flag^TTR secretion is observed. To better determine the relative amounts of secreted ^Flag^TTR for Hspa13 variant co-ovexpression, we immunoprecipitated ^Flag^TTR from the media using anti-Flag antibody. Immunoprecipitation allows the small amount of residual TTR secreted from Hspa13^T230A^ cells to be visualized. It also reveals that there is a small population of immature TTR secreted with Hspa13 overexpression. Brefeldin A is a small molecule that rapidly collapses the Golgi, and is often used to decipher whether a protein is secreted through the canonical secretory pathway (90–92). Brefeldin A treatment abolishes secretion of the mature TTR population, but does not decrease the immature population. This indicates that while the mature TTR is trafficked through the secretory pathway, the immature portion is bypassing the Golgi through another mechanism, such as necrosis (**Figure S6B**).

We further considered another secretory protein, Alpha-1 antitrypsin (A1AT). A1AT is a metastable N-glycosylated hepatically secreted protein (93, 94). Destabilizing mutations in A1AT lead to its aggregation or degradation in the cell, suppressing secretion and ultimately leading to diseases associated with A1AT serum deficiency and/or hepatic aggregation (95). We co-overexpressed Hspa13 variants with an A1AT construct with Flag situated C-terminal to the endogenous signal peptide. Hspa13 overexpression does not inhibit A1AT import into the ER, nor does it noticeably affect its secretion (**Figures S6D-F**). Hspa13^T230A^ overexpression, on the other hand, substantially decreases A1AT levels in the cell and its secretion. We also observed a small amount of non-glycosylated secreted A1AT with overexpression of both Hspa13 ATPase mutants (**Figure S6E**). The identity of the band was confirmed by treatment with PNGase F, which removes N-linked glycans and resolves both bands together (**Figure 6C**). Because glycan occupancy is linked to signal sequence identity, and weaker signal sequences show greater reliance on translocon-associated complexes such as TRAP and Sec62/63 (15, 96), we considered whether Hspa13 overexpression would have a larger effect on a protein harboring a weaker signal sequence. We replaced the signal sequence on ^Flag^A1AT with that of preprotrypsin (**Figure 6D**), which has a far shorter hydrophobic region; hydrophobicity is necessary to engage both SRP and Sec61 (97, 98). The A1AT sequons are at N70, N107, and N271 are far from the N-terminus (99, 100). As is the case for ss_A1AT_^Flag^A1AT, ss_ppt_^Flag^A1AT is secreted solely in the mature, glycosylated form (**Figure 6C**). However, the sensitivity to Hspa13 is now much greater, with Hspa13^WT^ overexpression now inducing noticeable secretion of immature, non-glycosylated ^Flag^A1AT, while Hspa13^T230A^ expression now leads to secretion of almost solely the non-glycosylated form. Hence, it is clear that in addition to its effects on ER translocation and cytosolic proteostasis, Hspa13 expression affects protein maturation in the ER.

### Hspa13 knockout impairs ER proteostasis

Having established that Hspa13 overexpression and mutation inhibit translocation, we looked to see the effect of ablating Hspa13. We generated *HSPA13*^*–/–*^ cells using the *Streptomyces pyogenes* CRISPR/Cas9 system (**Figure 7A** and **Figure S7A**) (101). Single clones were assessed for Hspa13 expression, and two *HSPA13*^*–/–*^ lines chosen for expansion: 1A3 and 12F4. While 12F4 grows normally and has similar BiP expression to HEK293T cells, 1A3 grows slowly and has highly upregulated BiP expression, suggesting constitutive UPR activation (**Figure 7B** and **Figure S7B**). In the 12F4 line, TTR maturation appears unaffected as compared to the parental line (**Figure 7B**). However, upon treatment with the canonical UPR chemical inducer thapsigargin (Tg), we observe a few differences between the *HSPA13*^*–/–*^ and parental lines. First, Tg induces high molecular weight and insoluble TTR aggregates in 12F4 *HSPA13*^*–/–*^ cells (**Figures 7B,C**). Second, while Tg induces immature TTR in both cell lines, presumably through the pre-emptive quality control pathway (44, 45), the effect is exacerbated in the 12F4 *HSPA13*^*–/–*^ cells. Finally, the sensitivity to Tg is greater in 12F4 *HSPA13*^*–/–*^ cells as determined by the resazurin metabolic activity assay (**Figure 7D**). The 1A3 *HSPA13*^*–/–*^ cell line, by contrast, is resistant to Tg toxicity (**Figure S7C**), consistent with its UPR activation. Hormetic UPR activation has been shown to both enhance TTR quality control and to protect against Tg toxicity (71, 78, 79, 102).

**Figure 7.**
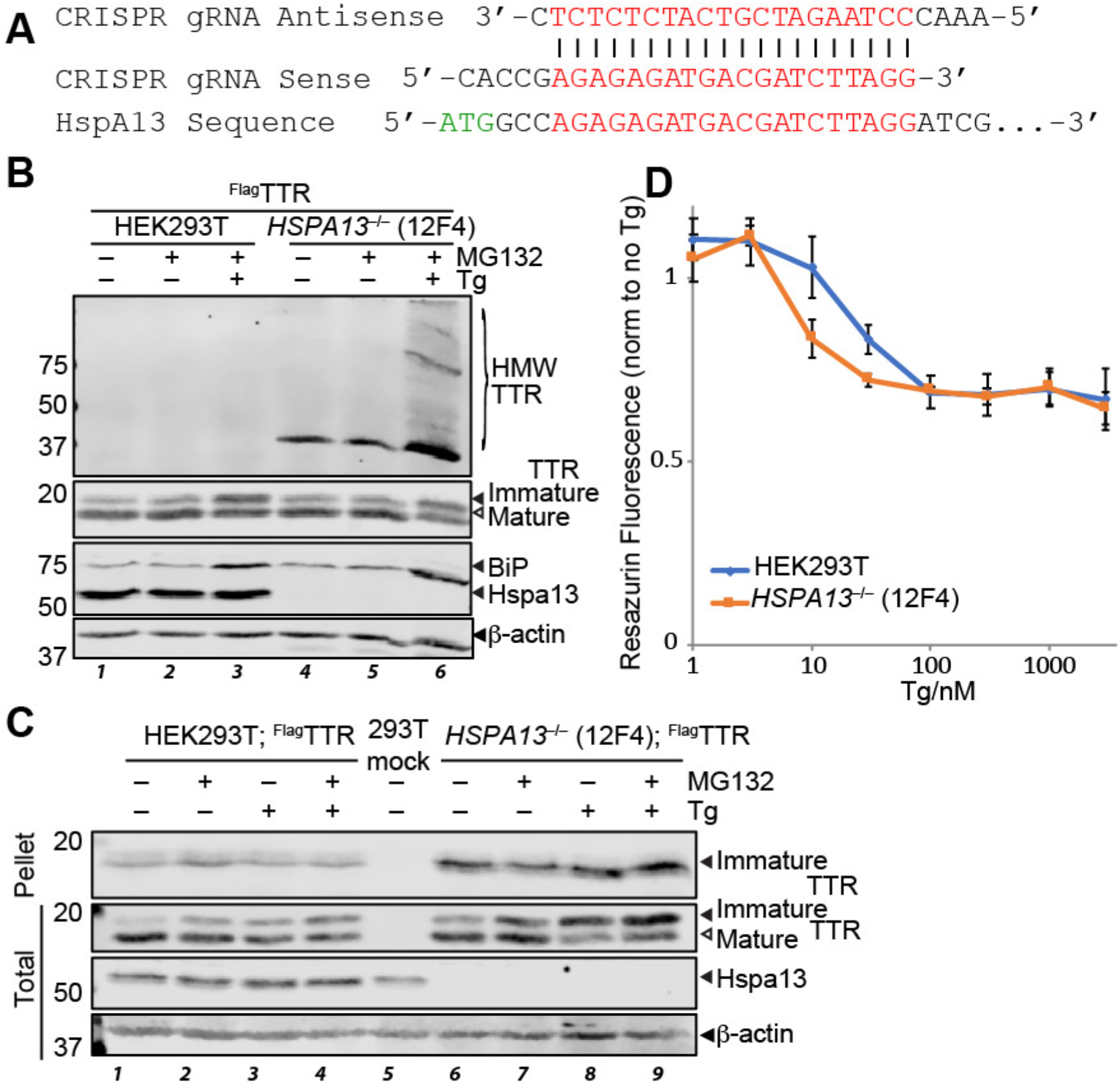
**A**. Schematic of gRNA sequences used to direct Cas9 cleavage to *HSPA13*. **B**. Representative immunoblot of SDS-PAGE separated lysates from HEK293T or *HSPA13*^*–/–*^ (12F4) cells overexpressing ^Flag^TTR, and treated with MG132 (100 nM; 16 h) and/or Tg (10 nM; 16 h) as indicated. HMW TTR is resolved on 4-20% gradient gels following 5 min. boiling in reducing Laemmli buffer, while immature and mature TTR monomers are resolved using 15% gels after 20 min. boiling. **C**. Representative immunoblot of SDS-PAGE separated lysates from HEK293T or *HSPA13*^*–/–*^ (12F4) cells overexpressing ^Flag^TTR, and treated with MG132 (200 nM; 16 h) and/or Tg (50 nM; 16 h) as indicated. Lysates were fractionated as described in **Figure 4C .D**. Normalized resazurin fluorescence (ex: 560 nm; em: 590 nm) signal of HEK293T or *HSPA13*^*–/–*^ (12F4) cells incubated with Tg (16 h) as indicated. Cells were seeded at 15,000 cells per well 2 h prior to Tg treatment (16 h). Media containing resazurin (25 μg/mL) was added to cells 2 h prior to measurement (n = 3).

## DISCUSSION

ER delivery of soluble secretory proteins is canonically encoded by the signal peptide. However, differences in signal peptide chemistry and cellular stress can influence both the targeting of proteins to the ER, as well as the maturation of proteins in the ER. Mislocalization and differential maturation in turn can affect both cytosolic and secretory proteostasis (4, 103–105). To understand the substrate-specific stringency of the translocon, as well as how stress can modulate this stringency, we need to determine the cellular components that modulate translocational activity. Herein, we have demonstrated that Hspa13 function impacts the stringency of translocational substrate import and maturation of two model secretory proteins: TTR and A1AT. Hspa13 specifically binds a majority of translocon subunits, as well as members of complexes associated with cotranslocational glycosylation and signal peptide cleavage. This is in sharp contrast to the other ER Hsp70, BiP, which has a wide-ranging interaction network that does not particularly feature translocon components (106, 107). In combination with previous reports of Hspa13 expression levels sensitizing mice to prion toxicity (21, 108), and to the synthetic toxicity of signal peptidase inhibition with *HSPA13* knockout (20), the present work further points to a role of Hspa13 in regulating the import and processing of nascent secretory proteins.

While these studies demonstrate that Hspa13 can interfere with secretory protein import and processing, they do not demonstrate that Hspa13 acts in such a manner in a particular disease state. TTR is not natively expressed in HEK293T cells, nor does it harbor a Flag tag after its signal sequence. Indeed, we see that TTR without the Flag tag is less prone to mislocalization, and the Hspa13 overexpression effect is severely decreased. Even in this case, the ATPase mutants promote mistargeting. Generally, every effect that we observe as a consequence of Hspa13^WT^ overexpression is greater for Hspa13^K100A^, and even greater for Hspa13^T230A^. Why would loss of ATPase activity exacerbate an effect associated with overexpression? An answer could be related to the function of the nucleotide binding domain (NBD) in full-length Hsp70s (109). The substrate binding domains in these proteins consist of two domains, SBDα and SBDβ. The ATP-bound form of Hsp70 forms rigid interactions between NBD and SBDβ. This interaction inhibits ATP hydrolysis, while also preventing tight binding to substrate through SBDα-SBDβ clamping. Hsp40 J-domain and substrate binding both serve to modulate this interaction, allowing the NBD to more readily access a conformation that is competent to hydrolyze ATP, thus releasing SBDβ. Hspa13 lacks the SBD, but does contain most NBD sites that interact with the Hsp40 J-domain. If this surface dysregulates import phenotypes through a direct binding event, then Hspa13 overexpression would increase the available Hspa13 for binding. In turn, ATPase mutants would maintain this surface in a higher-binding affinity conformation. We do see Hspa13 interactions with two translocon-associated Hsp40 proteins associated with post-translational ER import through Sec61: Sec62 and Sec63 (14). Hspa13 might interact with these proteins to affect their function, or indirectly compete with their association with BiP. Consistent with this possibility, A1AT with a weaker signal sequence was more sensitive to Hspa13 overexpression. Weaker signal sequences increase the reliance on post-translational translocation (15, 110). Further investigations will need to establish the molecular mechanism by which Hspa13 regulates import, as well as how this depends on the many other chaperones and stress factors that bind to and regulate the translocon (14, 111, 112), yielding a polydisperse distribution of translocon compositions (113, 114) that impact import and maturation (115).

Pre-emptive quality control is one of the least characterized consequences of UPR activation. Given that Hspa13 is a UPR-responsive target, and that it leads to TTR import inhibition, one might speculate that Hspa13 upregulation mediates pre-emptive quality control by the UPR. However, preemptive quality control is not interrupted in either of the *HSPA13*^*–/–*^ cell lines, establishing that Hspa13 does not mediate this process. Rather, Hspa13 knockdown aggravates the maturation (and presumably import) inhibition induced by thapsigargin in the 12F4 line. In the 1A3 clonal *HSPA13*^*–/–*^ line, where constitutive UPR activation offers hormetic protection, thapsigargin interferes with TTR maturation to a similar extent as the parental HEK293T line. In the absence of stress, *HSPA13* knockout also does not affect TTR maturation. Nevertheless, TTR aggregation is greater in the *HSPA13* knockout line, demonstrating impaired ER proteostasis. Similarly, Hspa13 overexpression leads to increased association between mature TTR and ER chaperones. It is clear that modulating Hspa13 levels in either direction is deleterious for the ER.

## EXPERIMENTAL PROCEDURES

### Plasmids

Expression constructs for Hspa13 and its variants were prepared in the pDEST30 vector. Hspa13 was amplified from HEK293T derived cDNA and ligated into the pENTR1A vector using *SalI* and *EcoRV* enzymes. HYOU1 was amplified from HEK293T-derived cDNA and ligated into the pENTR1A vector using *SalI* and *EcoRI* enzymes. The genes were then transferred into the pDEST30 vector by homologous recombination using the Gateway kit (Thermo). Hspa13 variants were prepared by site-directed mutagenesis. Hspa13^Flag^ was prepared from the Hspa13.pDEST30 plasmid by PIPES cloning (116). ERjd3 and BiP plasmids have been reported (88). ^Flag^TTR and ^Flag^TTR^D18G^ in the pcDNA3.1 vector have been reported (71). Ubiquitin^HA^ in the pcDNA backbone was a gift from Edward Yeh (Addgene plasmid # 18712 ; http://n2t.net/addgene:18712 ; RRID:Addgene_18712) (76). ^cyt^APEX2 in the pcDNA3 backbone was a gift from Alice Ting (Addgene plasmid # 49386 ; http://n2t.net/addgene:49386 ; RRID:Addgene_49386) (51). pCMV-^ER^HRP^N175S^ was a gift from Joshua Sanes (Addgene plasmid # 79909 ; http://n2t.net/addgene:79909 ; RRID:Addgene_79909) (50). CRISPR vector pX330A-1×2 was a gift from Takashi Yamamoto (Addgene plasmid # 58766 ; http://n2t.net/addgene:58766 ; RRID:Addgene_58766) (101). Targeting primers were ligated into BbsI-cleaved pX330A-1×2 to generate the *HSPA13*-targeting plasmid. Primers for all cloning procedures described above are listed in **Table S4**.

### Mammalian Cell Culture

HEK293T cells and derived lines were grown in DMEM (Corning) with 10% FBS (Seradigm), penicillin-streptomycin antibiotic cocktail and L-glutamine. SH-SY5Y cells were similarly grown in DMEM/Ham’s F-12 with the same supplements. Calcium phosphate transfection solution was performed as previously reported (54) with media change within 16 h of transfection. Unless otherwise indicated, cells were harvested by rinsing in 1x PBS (HyClone), scraping from the plates, and lysis in radioimmunoprecipitation buffer (RIPA; 50 mM Tris pH 7.4, 150 mM NaCl, 1% Triton X100, 0.5% sodium deoxycholate, 0.1% SDS) containing cOmplete protease inhibitor (Roche Diagnostics). Cells expressing Ub^HA^ were lysed in the presence of 12.5 mM N-ethyl maleimide. Lysis involved resuspension of cells and incubation on ice for 15 minutes, followed by a full-speed hard spin at 4°C for 20 minutes. Media conditioning was performed for 24 h, starting 12 to 16 h post-transfection, unless otherwise indicated. Conditioned media was collected and spun at 700 *x g* for 5 min to remove cellular debris. For Flag enrichments, M2-anti-Flag dynabeads (Sigma) were incubated with cultured media and rotated overnight at 4 °C. Protein was eluted by boiling in 4X Laemmli buffer, and then eluates collected and reduced by 20 mM DTT and further boiling for 20 minutes.

### Resazurin assays

Cells were incubated with DMEM containing resazurin (25 μg/mL; Acros Organics) for 2 hours. Fluorescence from cells was measured via a Synergy H1 microplate reader set for 560 nm excitation and 590 nm emission.

### Proximity Labeling

We used ^ER^HRP and ^cyt^APEX2 labeling to biotinylate proteins that are ER-lumenal space and cytosol targeted, respectively (46, 117, 118). Unless otherwise indicated, cells were treated with 1 μM MG132 treatment for 16 h prior to harvesting to inhibit proteasomal degradation. 30 min prior to harvest, the media was changed to fresh complete media containing 500 μM biotin phenol and 1 μM MG132. Immediately prior to harvesting, aqueous H_2_O_2_ (Fisher) was added to a final concentration of 1 mM, the plates were gently agitated, and incubated for exactly 1 min at ambient temperature. The reaction was quenched by washing three times with DPBS containing 5 mM Trolox, 10 mM sodium ascorbate and 10 mM sodium azide. Cells were then lysed in RIPA buffer supplemented with protease inhibitor cocktail, 5 mM Trolox, 10 mM sodium ascorbate and 10 mM sodium azide. Biotinylated protein was purified by incubation with avidin beads overnight with rotating at 4 °C. Beads were washed with the following buffers to remove nonspecific binders: twice with RIPA lysis buffer, once with 1.0 M KCl, once with 0.1 M Na_2_CO_3_, once with 2.0 M urea in 100 mM Tris, pH 8.0, and twice with RIPA lysis buffer. Biotinylated proteins were eluted by boiling the beads in elution buffer containing 12% SDS, 47% of glycerol in 60 mM Tris pH 6.8, with 10 mM DTT and 2 mM biotin for 10 min. The eluate was collected by centrifuging the beads and boiled for another 10 min.

### Ultracentrifugation

Normalized cell lysates were spun in a Beckman Coulter Optima Max-XP for 4 hours at 77,000 x g. The TLA-55 rotor was refrigerated at 4°C and kept under vacuum during the ultracentrifugation. Pellets were rinsed with RIPA and solubilized with 8 M urea over 4 days at 4°C, then diluted with 3 parts RIPA.

### Immunoblotting

Samples were boiled in 16.7 mM dithiothreitol (DTT)/Laemmli buffer for 20 min at 100 °C and resolved in homemade sodium dodecylsulfate polyacrylamide gel electrophoresis (SDS-PAGE) gels cast in Bio-Rad Mini-PROTEAN glass plate assemblies. Electrophoresis began at 65 V to allow sample stacking in the 4% acrylamide stacking gel, followed by 175 volts for resolution in the resolving gel. A gel gradient from 4-20% was used for resolving high-molecular weight ^Flag^TTR smears while 12% or 15% was used for resolving ^Flag^TTR monomers. Gradient gels were prepared by aspirating 4% activated gel solution into a pipette, an equal volume of 20% solution below it, and then aspirating three bubbles to generate the gradient, followed by careful dispensing into the gel cassette for polymeration. Protein was transferred to nitrocellulose membrane via semi-dry method (Bio-Rad Trans-Blot Turbo) for 1 h, using Towbin’s buffer as a saturant for the filter paper padding. Blotted membranes were assessed for quality with Ponceau stain, then blocked with 5% milk in Tris-buffered saline (TBS) for 40 min at ambient temperature. The blots were serially rinsed with Tris-buffered saline with 1:1000 Tween 20 (TBST) for 5 minutes per rinse, then incubated with primary antibody solution (primary antibody diluted in 5% bovine serum albumin (BSA), 0.1% sodium azide, TBS) for 2 hours. The blots were serially-rinsed and incubated with secondary antibody solution (1:20,000 Li-COR near-IR fluorophore-conjugated secondary antibody diluted in 5% milk, TBS) for 20-40 minutes and image on a Li-COR Fc.

### PNGase F treatmen

Lysates (20 μg) and media immunoprecipitates were diluted to 10 μL in 2X glycoprotein denaturing buffer (NEB), and boiled for 10 min. at 100 °C. 2 μL GlycoBuffer 2 10X (NEB), 2 μL NP-40, 1 μL PNGase F (NEB), and 5 μL water were added and the solution incubated for 37 °C for 1 h.

Knockout Cell Preparation. Knockout lines were prepared by transient transfection of HEK293T cells with pX330A-1×2 plasmid harboring the chosen guide RNAi, and transfect cells selected with Zeocin. After selection, individual cells were monoclonally expanded in the and Hspa13 protein expression characterized to identify *HSPA13*^*+/+*^, *HSPA13*^*–/+*^ and *HSPA13*^*–/–*^ lines.

### Sample Preparation for Mass Spectrometry

Protein samples were cleaned by methanol-chloroform precipitation. Dried protein pellets were reconstituted in 1% RAPigest (Waters or AOBIOUS) in 100 mM HEPES pH 8.0. Disulfide bonds were reduced in 10 mM TCEP for 30 min in the dark and alkylated with 5 mM iodoacetamide for 15 min in the dark at RT. Proteins were then digested with trypsin at 1% (w/w) trypsin at 37 °C overnight. For each analysis, a triplicate of GFP control and a triplicate of Hspa13 overexpression were labeled with 6-plex TMT reagents (Thermo). Reagents were dissolved in acetonitrile and added to peptides (1:1) dissolved in 100 mM HEPES pH 8.0 to a final acetonitrile concentration of 40% (v/v). The reactions were incubated for 1 hour at RT, then quenched with ammonium bicarbonate at a final concentration of 0.4%. The samples were combined and evaporated to 10 uL then brought to 200 uL by solvent A (5% acetonitrile, 0.1% formic acid). The pooled sample was acidified with 5% formic acid to pH 2. Samples were incubated at 37 °C and centrifuged at 21100 x g for a few cycles to remove Rapigest.

### LC-MS analysis

LC-MS experiments were performed on a Thermo LTQ Velos Pro equipped with an EASY-nLC 1000 nanoLC (Thermo). Buffer A is 5% aqueous acetonitrile, 0.1% formic acid. Buffer B is 80% aqueous acetonitrile, 0.1% formic acid. Peptides were initially loaded on a triphasic MudPIT trapping column and washed with buffer A. The columns were prepared from 100 μm I.D. fused silica (Molex) with a KASIL frit and loaded with 2.5 cm each of 5 μm Aqua C18 resin, Jupiter SCX resin, and 5 μm Aqua C18 resin (Phenomenex). After washing with buffer A and bumping from the first C18 phase to the SCX phase with an organic gradient, peptides were iteratively transferred to the second C18 phase with increasing injections of ammonium acetate (119). After each injection the MuDPIT column was washed with buffer A and then the peptides separated over a 25 cm analytical column (100 μm ID, 5 μm tip pulled on a Sutter Instruments T-2000 tip puller) packed with 3 μm C18 resin (Phenomenex) using a gradient from 7% to 55% buffer B at a flow rate of 500 nL/min. A voltage of 3.0 kV was applied for electrospray ionization. Dynamic exclusion was employed with 120 s exclusion and a 2.0 mass window. MS1 and MS2 were acquired in the Orbitrap with nominal resolving power of 7500 at 400 m/z. Isolation was performed in the linear ion trap in data-dependent mode on the top ten peaks following each MS1 with 1.0 isolation width. Fragmentation was performed in the HCD cell with stepped collision energies of 32%, 38%, and 44%. Raw spectra were converted to mzml files using MSConvert (120) for analysis by FragPipe. The MS/MS spectra obtained from HCD fragmentation in the Orbitrap were identified and TMT intensities integrated in FragPipe (121, 122), against the Uniprot 07/11/2021 human proteome release (longest entry for each protein group) with 20429 proteins plus reverse sequences and common contaminants, with a maximum peptide-level FDR of 0.01. Cysteine alkylation (+57.02146 Da) and TMT modification (+229.1629 on lysine and N-termini) were set as fixed modifications, and half-tryptic cleavages were allowed. Keratins, non-human contaminants, and immunoglobulins were filtered from proteins lists. For lysate and pellet experiments, integrated intensities at the protein for each channel were normalized to the median intensity of that channel at the peptide level, and then p-values were moderated (123). For immunoprecipitation, p-values were generated based on our previously reported bait-prey correlation method (54). Local false discovery rates (q-values) were assessed using Storey’s modification of the method of Benjamini and Hochberg (55, 56). The Storey π_0_ factor was 1.0 for lysate and pellet experiments, and 0.13 for the AP-MS experiment.

## ACKNOWLEDGEMENTS

We thank E. Yeh for the Ub^HA^ plasmid, A. Ting for the ^cyt^APEX2 plasmid, J. Sanes for the ^ER^HRP^N175S^ plasmid, and T. Yamamoto for the pX330A-1×2 plasmid. Support was provided by the University of California.

## DATA AVAILABILITY STATEMENT

Mass spectrometry-based proteomics raw data and search results are available at the PRIDE Archive at PXD033498. Quantitation is available in Supporting Information **Tables S1-S3**.

