## Supplemental Figures and Supplemental Table S4 for "Hspa13 Regulates Endoplasmic Reticulum and Cytosolic Proteostasis Through Modulation of Protein Translocation"


| Page | Contents |
|---|---|
| S1 | **Table of Contents** |
| S2 | **Supplemental Figures** |
| S11 | **Table S4** |


**Tables S1-S3** are provided as an .xlsx file in **Supporting Information**.

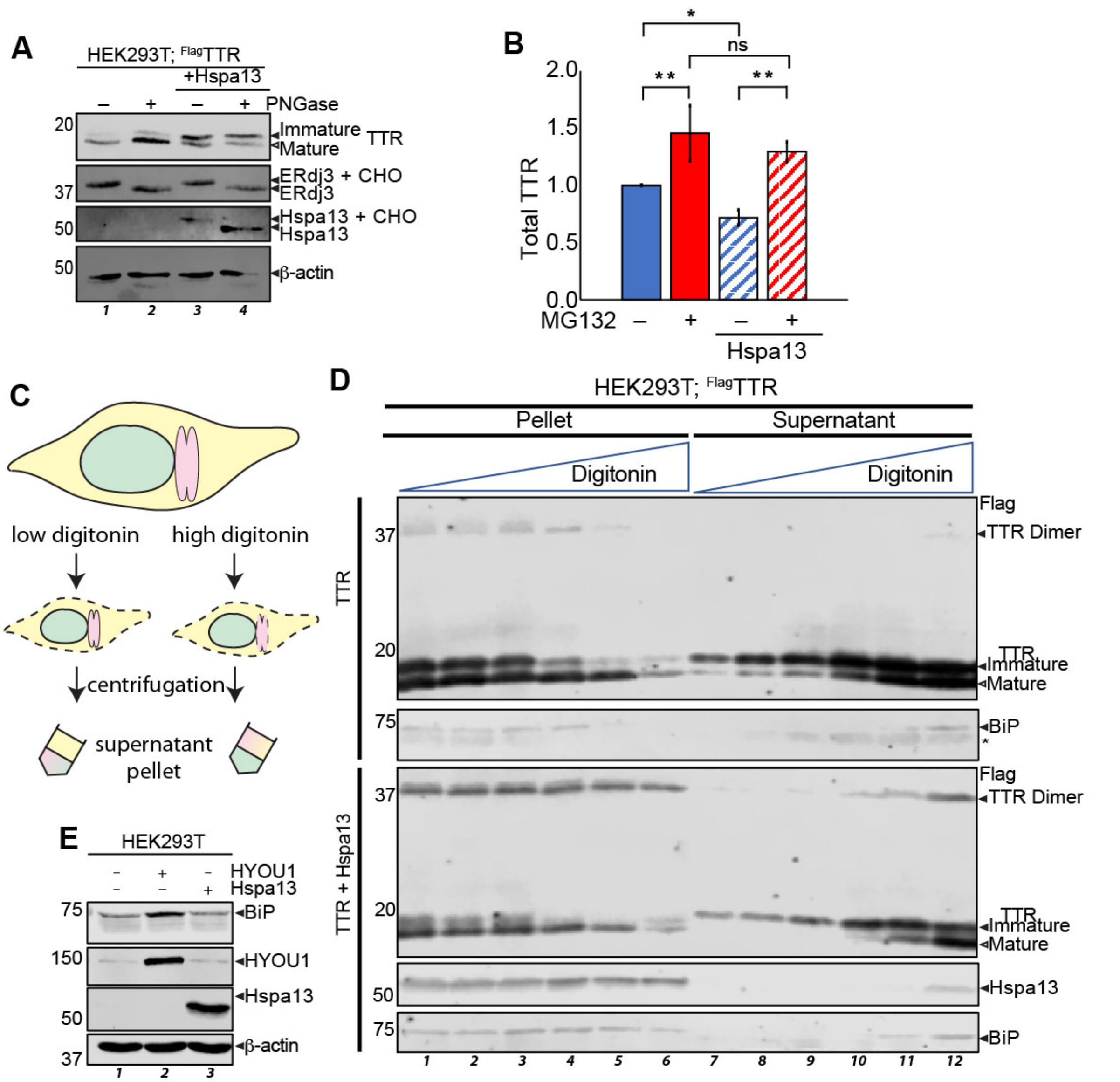


**Figure S1. A.** Immunoblot demonstrating lack of effect of PNGase F on $^{Flag}$TTR band structure, using SDS-PAGE separated lysates from HEK293T cells overexpressing $^{Flag}$TTR and/or Hspa13 as indicated. Removal of known glycans on Erdj3 and Hspa13 serve as a positive control. **B**. Densitometric quantification of total monomeric $^{Flag}$TTR from immunoblots such as in **Figure 1B**. Normalized to –Hspa13, –MG132. Error bars represent standard deviation (n = 5), single-factor ANOVA: $F = 14.0 > F_{crit} = 3.2$. Tukey's post-hoc HSD ** $p < 0.01$. **C**. Schematic of digitonin-based isolation of intact ER from HEK293T cells. **D**. Immunoblot of SDS-PAGE separated lysates and resolubilized pellets from HEK293T cells overexpressing $^{Flag}$TTR and Hspa13 as indicated and treated with gradient concentration of digitonin at 0, 0.005, 0.01, 0.05, 0.1, 0.25 % for 30 min. on ice, prior to 10,000 *x g* for 10 min. Pellets were resuspended in 1% Triton, 20 mM HEPES, pH 7.5, 100 mM NaCl, 1 mM EDTA, 1% Triton X100 for 10 min., and then boiled in reducing Laemmli buffer * indicates a band from a previous round of blotting. **E**. Immunoblot of SDS-PAGE separated HEK293T lysates overexpressing Hspa13 or HYOU1 as indicated.

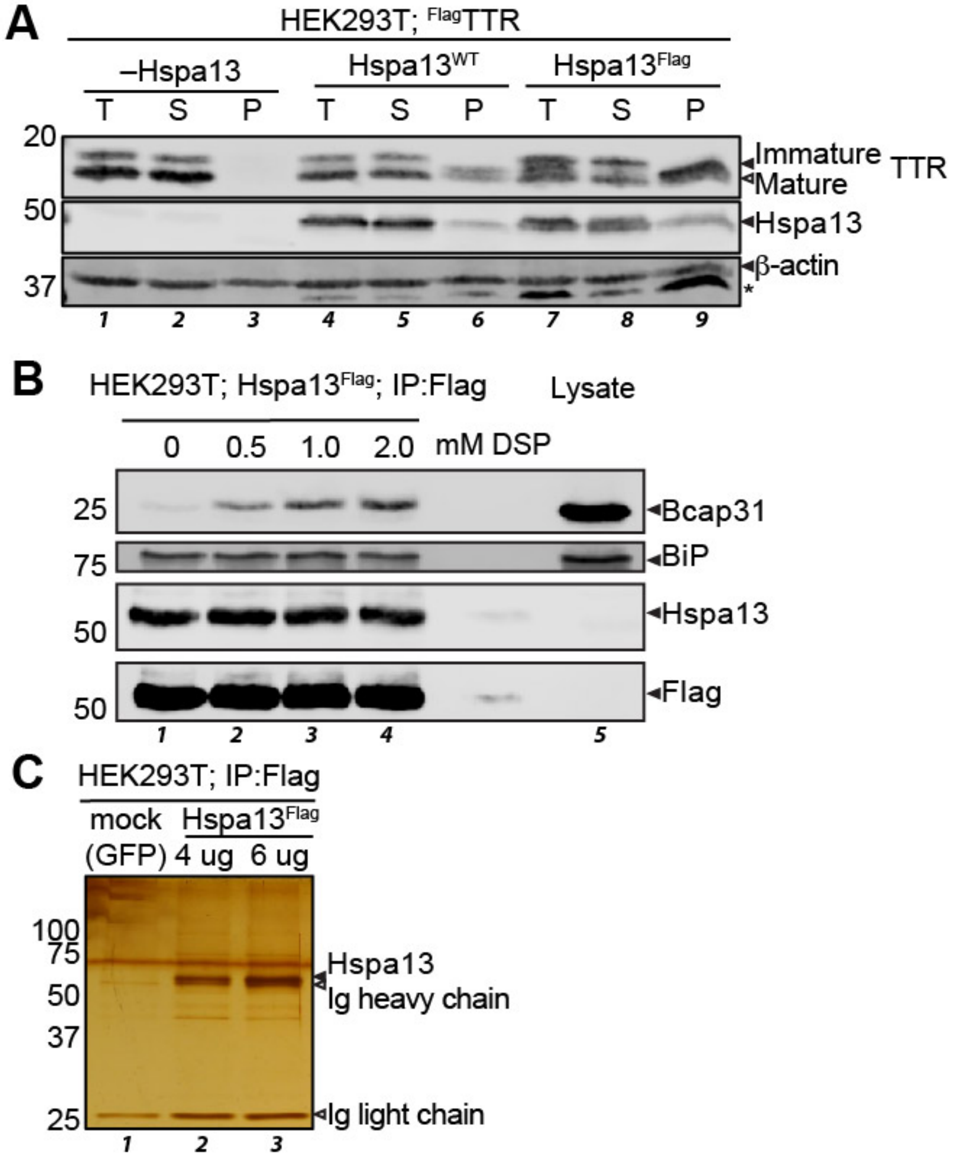


**Figure S2. A**. Immunoblot of SDS-PAGE separated lysates from HEK239T cells overexpressing $^{Flag}$TTR and Hspa13 or Hspa13$^{Flag}$ as indicated. Pellets and Supernatants were extracted as described in **Figure 4C**. The pellets in this experiment were enriched the intermediate fraction more than the immature fraction, in contrast to the rest of our replicates. This might have been due to an ultracentrifuge failure that arrested the spin prior to the full 4 h. **B**. Immunoblot of SDS-PAGE separated M2 anti-Flag immunoprecipitates from lysates of HEK293T cells expressing Hspa13$^{Flag}$. **C**. Silver stain of M2 anti-

FLAG immunoprecipitates form lysates of HEK239T cells expressing Hspa13$^{Flag}$ with the indicated amount of transfected plasmid.

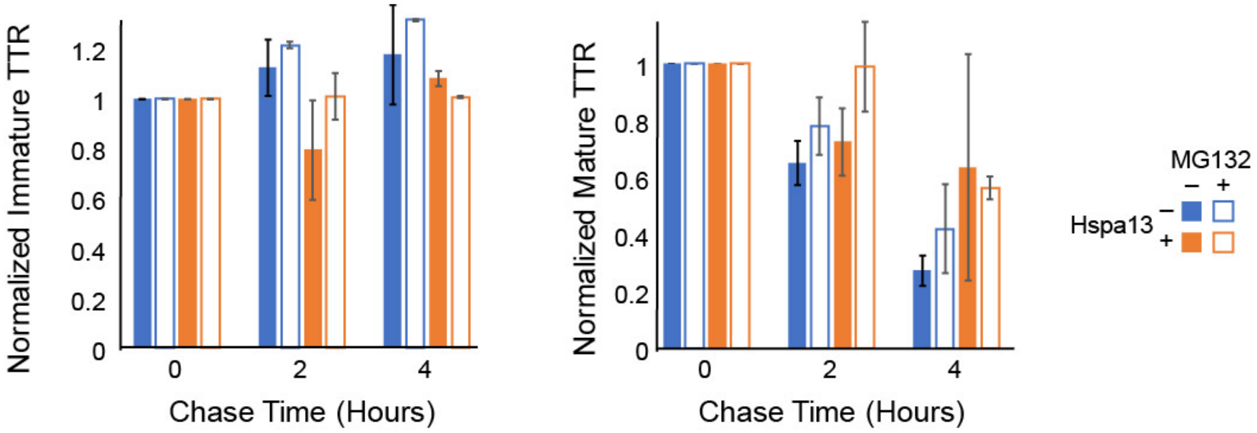


**Figure S3.** Quantifications of Immature and Mature TTR fractions respectively from **Figure 3A**, normalized to band densitometries at the beginning of the CHX chase. Error bars represent standard deviation (n = 2).

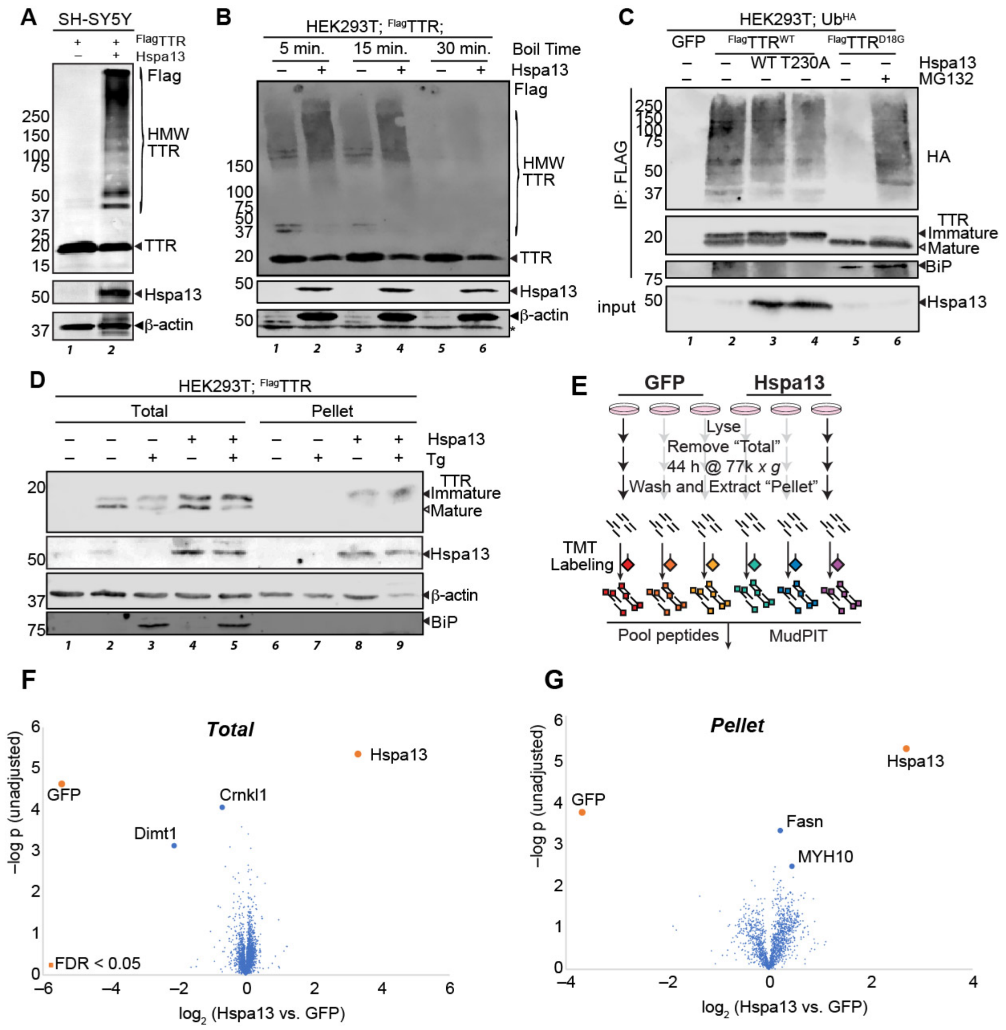

**Figure S4. A.** Immunoblot of SDS-PAGE separated lysates from SH-SY5Y cells overexpressing $^{Flag}$TTR and Hspa13 as indicated. **B**. Immunoblot of SDS-PAGE separated lysates from HEK293T cells overexpressing $^{Flag}$TTR and Hspa13 as indicated, with varying boiling times for the lysates. **C.** Representative immunoblot of SDS-PAGE separates M2 anti-Flag immunoprecipitates from HEK293T cells overexpressing indicated gene products. MG132 treatment, where indicated, was for 16 h at 1 µM. Cells were lysed in the presence of 12 mM N-ethylmaleimide. **D**. Representative immunoblot of SDS-PAGE separated lysates and pellets from HEK293T cells expressing $^{Flag}$TTR or Hspa13 as indicated and separated as described in **Figure 4C**. Tg treatment was 100 nM for 16 h. **E**. Schematic describing preparation of samples for mass spectrometry-based proteomics of lysates and pellets. **F**. Volcano plot illustrating differences in protein abundances in lysates from HEK293T cells overexpressing Hspa13 vs. mock (GFP). Only Hspa13 and GFP are significantly below a 5% FDR threshold. **G**. Volcano plot illustrating differences in protein abundances in the insoluble fractions from HEK293T cells overexpressing Hspa13 vs. mock (GFP). Only Hspa13 and GFP are significantly below a 5% FDR threshold.

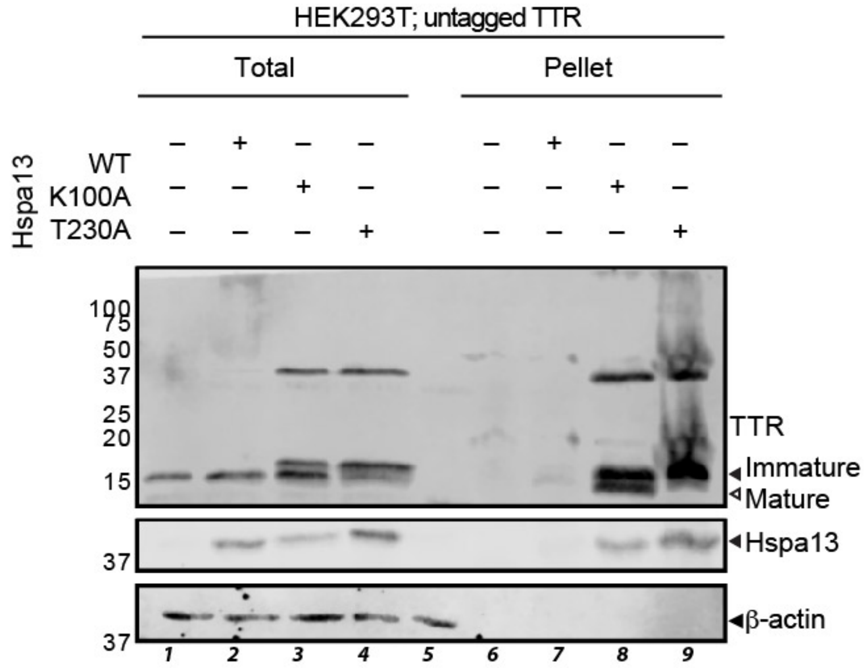


**Figures S5**. Representative immunoblot of SDS-PAGE separated lysates and pellets from HEK293T cells expressing untagged TTR and Hspa13 as indicated and separated as described in **Figure 4C**. The pellets are each 18-fold concentrated as opposed to the total lysate, and the middle lane is a naive lysate sample. The TTR band intensity in lane 7 (Hspa13$^{WT}$, pellet) is faint, but was similar in all four replicates.

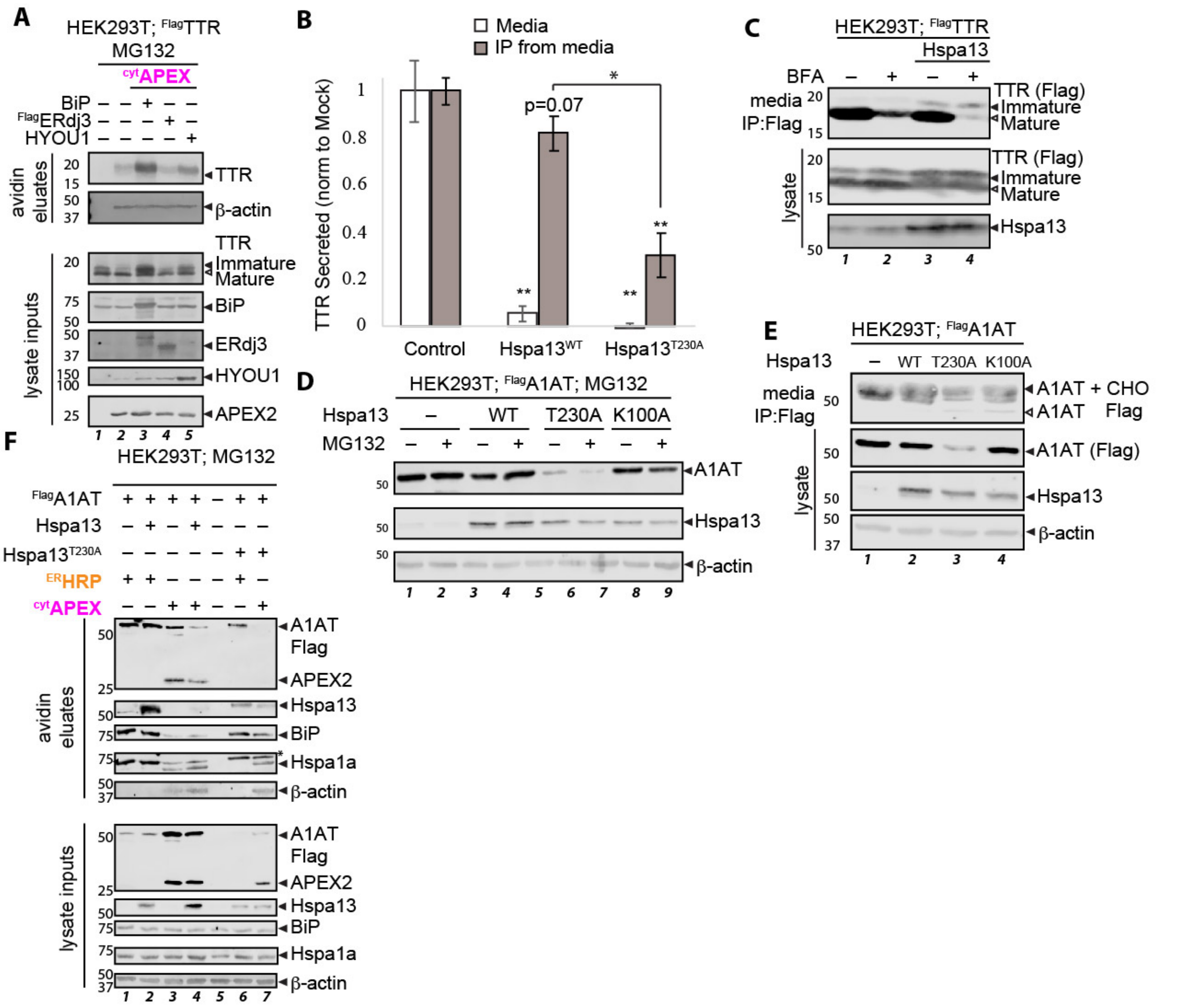


**Figure S6. A.** Representative immunoblot of SDS-PAGE separated lysates from HEK293T cells overexpressing $^{Flag}$TTR and major ER chaperones as indicated, and either $^{ER}$HRP or $^{cyt}$APEX. Cells were treated with BP (30 min. pre-harvest) and $H_2O_2$ (1 min. pre-harvest), quenched, and the biotinylated protein purified from lysate using avidin beads. All cells were treated with 1 µM MG132 for 16 h prior to lysis. Full blot images are provided in the Supplemental Gel Images File. **B**. Quantification of TTR levels in the media and media IP from **Figure 6B**. Error bars represent standard deviation (n = 3). For media, single-factor ANOVA $F = 151.8 > F_{crit} = 5.1$. For media IP, $F = 65.6 > F_{crit} = 5.1$. * represents $p < 0.001$, ** represents $p < 0.0001$. Significance markers unless otherwise indicated are with reference to Mock. Although comparisons of media input and immunodepleted media show quantitative depletion of $^{Flag}$TTR across conditions, there appears to be some saturation of signal with immunoprecipitation. **C**. Representative immunoblot of SDS-PAGE separated lysates and conditioned media M2 anti-Flag immunoprecipitates from HEK293T cells overexpressing $^{Flag}$TTR and Hspa13 as indicated, and treated with brefeldin A (BFA). The cells were treated with BFA (100 ng/mL) for 1 h, then the media replaced with fresh BFA-containing media for 6 h conditions, followed by IP. **D,E.** Representative immublots of SDS-PAGE separated lysates and media immunoprecipitates from HEK293T cells overexpressing indicated gene products. MG132 treatment, where indicated, was for 16 h at 1 µM. **F**. Representative immunoblot of SDS-PAGE separated lysates from HEK293T cells overexpressing $^{Flag}$A1AT and Hspa13 variants as indicated, and either $^{ER}$HRP or $^{cyt}$APEX. Cells were treated with BP (30 min. pre-harvest) and $H_2O_2$ (1 min. pre-harvest), quenched, and the biotinylated protein purified from lysate using avidin beads. All cells were treated with 1 μM MG132 for 16 h prior to lysis.

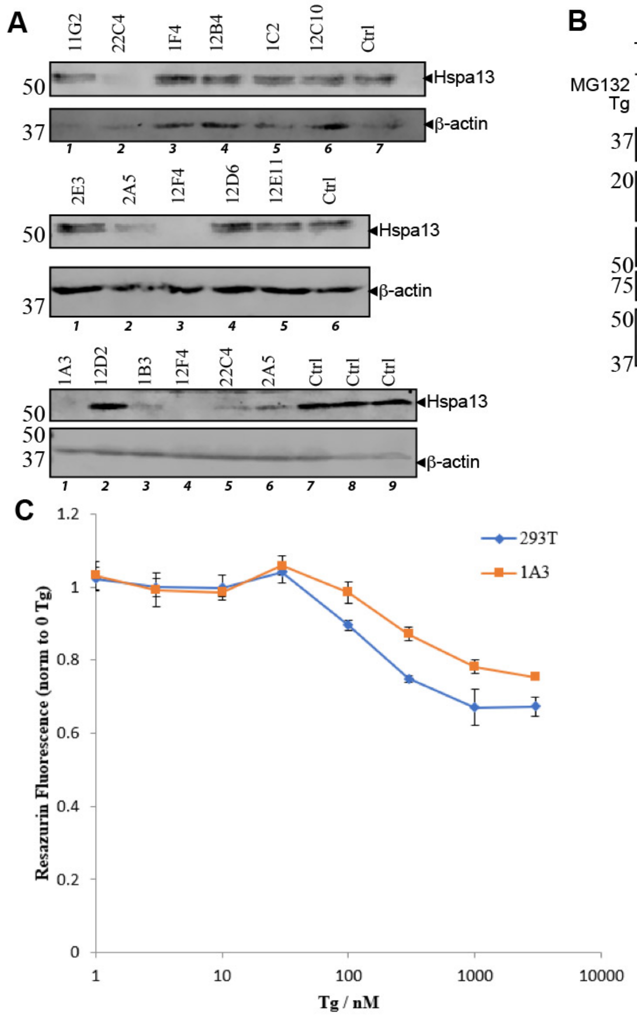


**Figure S7**. **A**. Immunoblot of SDS-PAGE separated lysates from monoclonal HEK293T lines stably transfected with *HSPA13*-targeting CRISPR vector. **B**. Immunoblot of SDS-PAGE separated lysates from *HSPA13*$^{-/-}$ (1A3) or HEK293T cells overexpression $^{Flag}$TTR and treated with MG132 (100 nM, 16 h) or Tg (10 nM, 16 h). **C**. Normalized (to no Tg) fluorescence signal of 1A3 vs 293T cells in the same well-plate, incubated with indicated concentrations of Tg for 16 h. Cells received a media change to media with resazurin (25 µg/mL) 135 min prior to fluorescence measurement (560 nm excitation, 590 nm emission). Error bars represent standard error of the mean (n = 3).

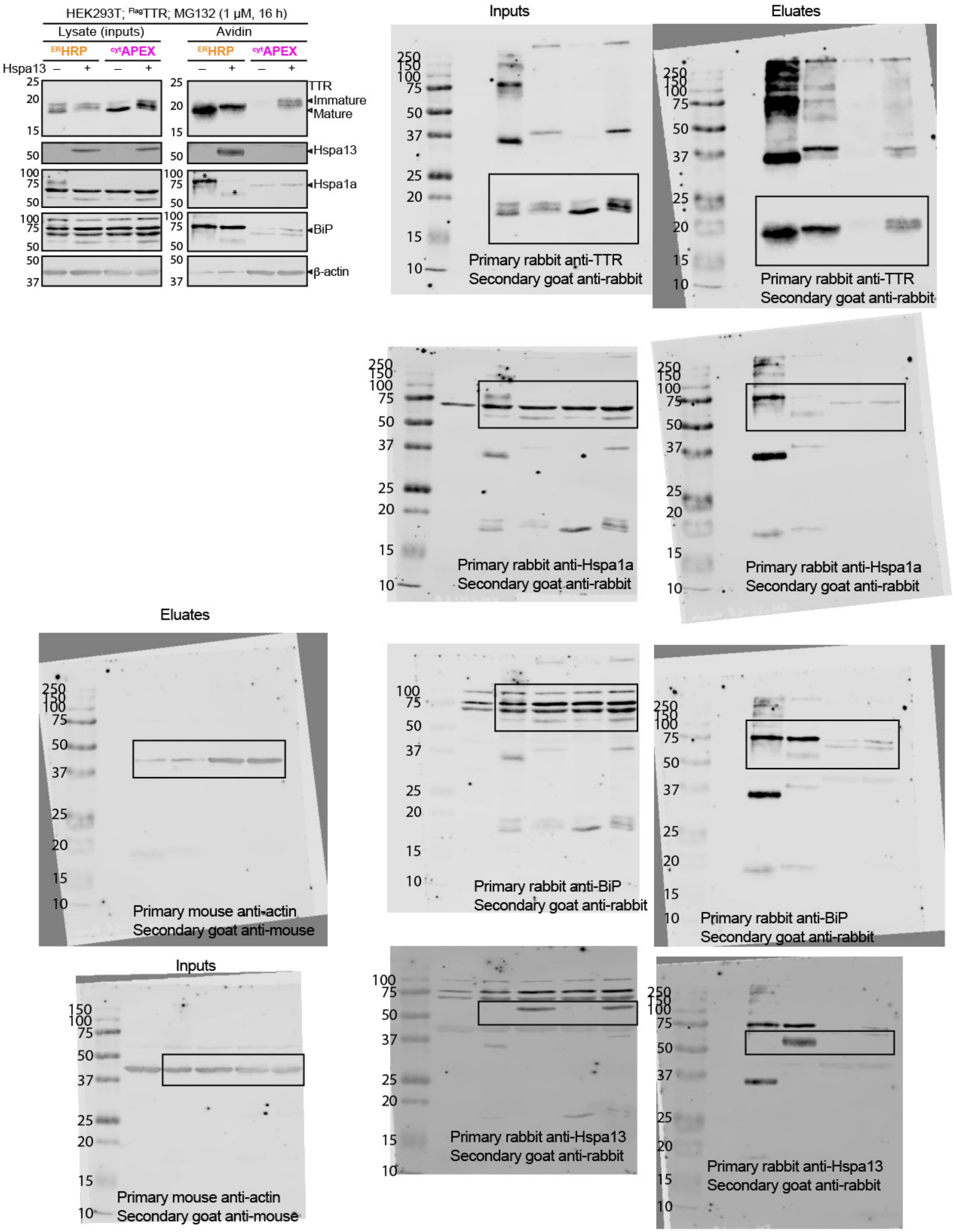
HEK293T; FlagTTR; MG132 (1 μM, 16 h)
Lysate (inputs)
Avidin
ERHRP
cytAPEX
Hspa13
TTR
Immature
Mature
Hspa13
Hspa1a
BiP
β-actin
Inputs
Eluates
Primary rabbit anti-TTR
Secondary goat anti-rabbit
Primary rabbit anti-Hspa1a
Secondary goat anti-rabbit
Primary mouse anti-actin
Secondary goat anti-mouse
Primary rabbit anti-BiP
Secondary goat anti-rabbit
Primary rabbit anti-Hspa13
Secondary goat anti-rabbit

**Figure S8**. Full blot images of **Figure S1E**, to show interference of TTR blotting. $^{ER}$HRP heavily oxidizes TTR, leading to the formation of detergent-resistant aggregates (Ref. 46 of the main text, Lyu et al). $^{Cyt}$APEX does not have the same effect; rather the high molecular weight TTR seen with Hspa13 overexpression is a consequence of Hspa13 overexpression and not labeling (**Figure 4A**).

### Table S4: Primers/Oligonucleotides

| **Name/Use** | **Sequence (5' → 3')** |
|---|---|
| HYOU1 Forward | AAC ATA GTC GAC TAT GGC AGA CAA AGT TAG GAG G |
| HYOU1 Reverse | ATG TTG AAT TCT TAT AGT TCG TCG TTC TTC AAA GGC C |
| HSPA13 Forward | AAC ATA GTC GAC TGT GAT GGC CAG AGA GAT G |
| HSPA13 Reverse | ATG TTG ATA TCA GTT GAA GTT GGT TTT TTG TAA ATG CTT A |
| HSPA13 K100A SDM Forward | CAC AAT ATA TGA TGC CGC AAG ATT CAT AGG CAA GAT TTT TAC CGC AGA GTT GGA GGC |
| HSPA13 K100A SDM Reverse | GAA TCT TGC GGC ATC ATA TAT TGT GTT TTG AGG ATT TGA ATC TGC CAG CTC TAC GCT TTC |
| HSPA13 T230A SDM Forward | GCG GAG GAG CTC TAG ATG TGT CTT TAC TGA ATA AAC AAG GAG GG |
| HSPA13 T230A SDM Reverse | CAT CTA GAG CTC CTC CGC CCA AGT CTA TCA CCA AGA CGT GGA AG |
| HSPA13-FLAG Forward (cDNA) | GAC TAC AA GAC GAT GAC GAC AAG TAA CTA GAC CCA GCT TTC TTG TAC |
| HSPA13-FLAG Reverse (cDNA) | TTA CTT GTC GTC ATC GTC TTT GTA GTC GTT GAA GTT GGT TTT TTG TAA ATG C |
| HSPA13-FLAG Forward (vector) | GAC TAC AAA GAC GAT GAC GAC AAG TAA CTA GAC CCA GCT TTC TTG TAC AAA G |
| HSPA13-FLAG Reverse (vector) | TTA CTT GTC ATC GTC TTT GTA GTC GTT GAA GTT GGT TTT TTG TAA ATG CTT ATT G |
| HSPA13 CRISPR Sense | CAC CGA GAG AGA TGA CGA TCT TAG G |
| HSPA13 CRISPR Antisense | AAA CCC TAA GAT CGT CAT CTC TCT C |
